# Glutamine-driven mTORC1 activity enforces glycolytic bias, hyperactivation and aberrant proliferation in T cells in chronic B-cell leukemia

**DOI:** 10.64898/2026.06.09.731122

**Authors:** Nienke B Goedhart, Elena Camerini, Gaspard Cretenet, Anita E Grootemaat, Kirill Vlaskov, Chaja F Jacobs, Yannick d’Hargues, Nicole N van der Wel, Frank Vrieling, Bauke Schomakers, Michel van Weeghel, Kimberly M Bonger, Mark-David Levin, Eric Eldering, Arnon P Kater, Helga Simon-Molas

## Abstract

T-cell metabolic dysfunction is increasingly recognized as a hallmark of poor antitumor responses. Still, how distinct metabolic states shape T-cell function and which signaling pathways sustain them remains unclear. Here, we show that T cells from patients with chronic lymphocytic leukemia (CLL) adopt a hyperactivated phenotype characterized by high cytokine production, aberrant proliferation and a bias toward glycolysis, supported by sustained glutamine-driven mTORC1 activity. In parallel, T cells from these patients display alterations in mitochondrial network and cristae architecture, which further limit oxidative phosphorylation. mTORC1 inhibition reduces glucose dependence, restores mitochondrial metabolic engagement and normalizes T-cell activation and proliferation.

Together, these findings identify metabolic disbalance as a central contributor to the hyperactivated state of T cells in CLL. We propose a model in which reduced OXPHOS reflects both mitochondrial defects that pre-exist in T cells from patients before TCR engagement, and a failure to engage mitochondrial metabolism upon activation supported by the actionable target mTORC1.

**Highlights:**

- T cells from patients with chronic lymphocytic leukemia display high cytokine production and excessive division cycles upon CD3/28 engagement.
- mTORC1-sustained glycolysis and defects in mitochondrial structure compromise OXPHOS in hyperactivated T cells.
- Exogenous glutamine uptake sustains non-lysosomal mTORC1 activity in T cells.

## Introduction

Cellular metabolism is a central regulator of T-cell function and fate^1–3^. Upon activation, T cells undergo metabolic reprogramming characterized by increased aerobic glycolysis, often referred to as the glycolytic switch, which provides rapid energy and biosynthetic precursors to support proliferation and cytokine production^4,5^. While glycolysis has long been recognized as essential for T-cell activation, the importance of mitochondrial metabolism has more recently emerged as a key determinant of effective T-cell responses. Beyond supporting activation, mitochondrial metabolism is critical for long-term T-cell fitness, survival and memory formation^6–8^. Maintaining a balance between these metabolic programs is therefore essential to meet the energetic and biosynthetic demands of both acute and durable immune responses.

In cancer, T cells frequently fail to maintain this balance. Most tumor-infiltrating lymphocytes (TILs) exhibit impaired metabolic flexibility with defects in both glycolytic and mitochondrial metabolism, which can limit effector function, promote dysfunctional differentiation states and ultimately lead to T-cell exhaustion^9–11^. Several mechanisms have been identified as drivers of this global metabolic dysfunction in TILs. For example, constraints within the tumor microenvironment, such as nutrient and oxygen depletion, can restrict the resources available to T cells, limiting metabolic pathway activity^12,13^. Additionally, alterations in metabolic signaling, such as chronic AKT activation, can limit mitochondrial biogenesis, thereby reducing metabolic flexibility and driving T-cell exhaustion^11^. Together, these models suggest that both extrinsic nutrient limitation and intrinsic signaling alterations constrain T-cell metabolic flexibility in tumors, although this framework does not fully explain the spectrum of T-cell dysfunction observed across malignancies.

In chronic lymphocytic leukemia (CLL), T cells are consistently exposed to cancer cells for years due to the slow, progressive expansion of clonal CD5⁺ mature B cells that circulate between lymph nodes and peripheral blood. T cells from patients with CLL (hereafter, CLL T cells) display features resembling exhaustion, including skewing towards highly differentiated effector phenotypes and expression of inhibitory receptors^14,15^. However, unlike exhausted TILs, CLL T cells retain substantial capacity to produce cytokines, indicating that their functional state is distinct from classical T-cell exhaustion^16,17^. Yet, CLL T cells are not fully functional, as evidenced by their limited expansion capacity upon repetitive stimulation^18^ and the poor efficacy of chimeric antigen receptor (CAR)-T cell therapy in this disease^19^.

We and others have shown that functional defects in CLL T cells are accompanied by metabolic alterations, including reduced glucose metabolism in patients with impaired T-cell responses to in vitro stimulation, and increased production of mitochondrial reactive oxygen species (ROS)^20,21^. Recently, we demonstrated that these metabolic abnormalities worsen with disease progression and are associated with terminal differentiation and reduced CAR-T efficacy^17^. However, how tumor-imposed metabolic states shape T-cell function, and which signaling pathways drive these programs, remain poorly defined. By integrating analyses of T-cell phenotype, metabolism, and signaling downstream of CD3/CD28 engagement in CLL T cells, we address these questions and provide mechanistic insight into their regulation.

## Results

### T cells from CLL patients display excessive activation at late stages after CD3/28 engagement

In order to profile the activation response of T cells from CLL patients in the presence of their autologous malignant cells, we stimulated PBMCs from CLL patients with anti-CD3/CD28 antibodies for two-five days. Aged-matched healthy donors (HD) were used as controls. While some patients showed reduced expression of T-cell activation markers at day two post-stimulation (i.e., CD25, cytokines, CD107a, GZMB), others exhibited responses comparable to HD (Fig. 1A–B). CD25 expression correlated with other activation markers (CD71, CD69), supporting its use as a surrogate marker for activation (Fig. S1A). Although prior studies focused on hyporesponsive CLL T cells (patients 4–8, Fig. S1A), a subset of patients analyzed displayed preserved activation capacity (patients 1–3, Fig. S1A). Analysis of a larger independent cohort (N=37) confirmed this heterogeneity, with 43% of patients showing high activation and 57% showing poor responses (cutoff at 40% CD25+ in the CD4+ compartment, based on median 38,5% CD25+ CD4+ cells in this patient cohort; Fig. S1B). Similar heterogeneity was observed in the CD8+ compartment, with lower median percentage of CD25 expression (29,80%). Longitudinal sampling from CLL patients included in the B-cell malignancies biobank at Amsterdam UMC showed these responses were stable within individuals at different timepoints (T1-T3) (Fig. S1C). This stratification was not explained by immunological parameters (Fig. S1D-E), CMV status (Fig. S1F), or clinical prognostic markers of CLL (Fig. S1G–H), indicating that T-cell responsiveness to anti-CD3/CD28 stimulation is a consistent, patient-intrinsic trait.

**Figure 1.**
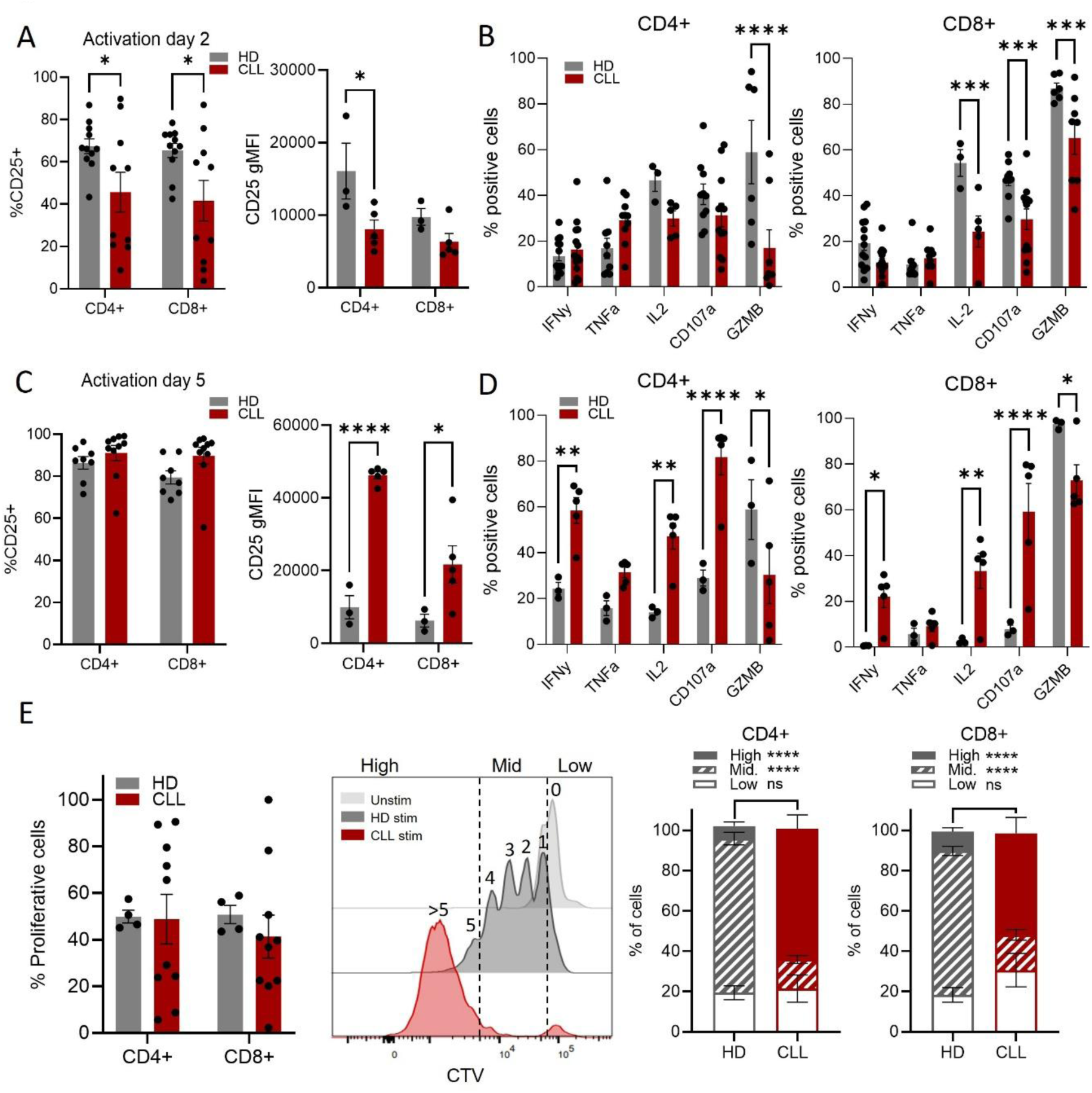
T cells from CLL patients display excessive activation at late stages after CD3/28 engagement. PBMCs from CLL patients or age-matched HDs were stimulated with αCD3/CD28 antibodies for five days, and flow cytometry was performed at day two (A–B) and day five (C–E). (A) Percentage of CD25⁺ T cells and CD25 expression levels. (B) Percentage of cytokine-positive and degranulating (CD107a⁺) T cells, determined by intracellular cytokine staining and surface CD107a staining. (C) Percentage of CD25⁺ T cells and CD25 expression levels. (D) Percentage of cytokine-positive and degranulating (CD107a⁺) T cells. (E) Representative proliferation profile of CD4⁺ T cells measured by CellTrace Violet (CTV) and example of the gating strategy. Quantification of the percentage of cells that entered division (% proliferating cells) and distribution of cells undergoing low (0–1 divisions), intermediate (2–4 divisions), or high (>5 divisions) proliferation (HD: N = 4, CLL: N = 10). Differences were analyzed using two-way ANOVA. A p-value < 0.05 was considered statistically significant. *p < 0.05; **p < 0.01; ***p < 0.001; ****p < 0.0001. Error bars represent mean ± SEM.

Despite the heterogeneous response observed across CLL patients at early stages of T-cell activation, by day five post-stimulation all patients displayed high activation levels. While the percentage of CD25⁺ T cells was comparable to HD (Fig. 1C), levels of CD25 and CD71 were significantly increased in CLL patients (Fig. 1C; Fig. S1I). The levels of CD25 and CD71 in CLL were also higher than those registered in HD T cells at day 2 (Fig. S1J), indicating not only a delay but a significantly different activation response in the T cells from patients at late stages after anti-CD3/CD28 stimulation. Consistent with an increased activation state, cytokine production and degranulation were elevated in T cells from CLL patients. In both CD4+ and CD8+ cells, the fraction of cells with intracellular production of IFNγ and IL-2 was increased, as well as surface levels of CD107a (Fig. 1D). This high activation phenotype (hereafter referred to as ‘hyperactivation’) was associated with altered proliferation dynamics. Although the fraction of proliferative cells was heterogeneous across CLL samples with some patients showing T-cell proliferation higher than HD and others lower (Fig. 1E), CellTrace Violet (CTV) profiling revealed a significant increase in highly proliferative T cells (>5 divisions) in PBMCs from all CLL patients analyzed. In contrast, T cells from HD showed a more balanced CTV profile with most cells showing mid-proliferation (2–4 divisions) (Fig. 1E). Thus, while early activation responses of CLL T cells to anti-CD3/CD28 stimulation are heterogeneous, all patients ultimately display features of a hyperactivated state with excessive division cycles. These findings prompted us to investigate to what extent this phenotype is supported by specific metabolic programming.

### Persistent glycolysis and impaired mitochondrial activity characterize activated CLL T cells

To characterize the metabolic profile of CD4+ and CD8+ T cells during activation, we applied CENCAT, a single-cell approach that uses protein synthesis as a surrogate for metabolic activity^22,23^ (Fig. S2A). Total viable CD4+ and CD8+ T cells were gated for these analyses. HD T cells showed higher dependence on glucose and higher glycolytic capacity when mitochondrial ATP was inhibited compared to unstimulated cells, consistent with a stimulation-induced glycolytic switch. Accordingly, dependence on mitochondrial ATP production and capacity to oxidize fatty acids (FA) and amino acids (AA) when glucose uptake was blocked were lower after stimulation (Fig. S2B). In CLL, two phenotypes were observed. In patients with preserved T-cell activation capacity (CD25+ ≥ 40%), CD4+ and CD8+ T cells showed lower mitochondrial dependence and higher glycolytic capacity than HD (Fig. 2A). This indicated a preference for glycolysis-derived ATP production over mitochondrial substrates in CLL T cells. In patients with poor activation responses, mitochondrial dependence was higher and glycolytic capacity was lower than in HD in CD4+ T cells, resembling the phenotype of unstimulated cells (Fig. S2C). No differences were observed in CD8+ T cells in this patient subgroup. At a later timepoint (day six post-stimulation), all CLL patients displayed significantly lower dependence on mitochondrial ATP production and lower capacity to oxidize FA and AA when glucose uptake was inhibited (Fig. 2B). Accordingly, patients showed higher glucose dependence and glycolytic capacity than HD T cells in both CD4⁺ and CD8⁺ compartments (Fig. 2B). This pointed to a skewing toward a glycolytic phenotype during T cell activation in CLL. In fact, while FA/AAO capacity significantly increased in HD T cells from days two to six, it declined in CLL T cells alongside increased glucose dependence (Fig. 2C), consistent with impaired switch towards mitochondrial oxidative metabolism.

**Figure 2.**
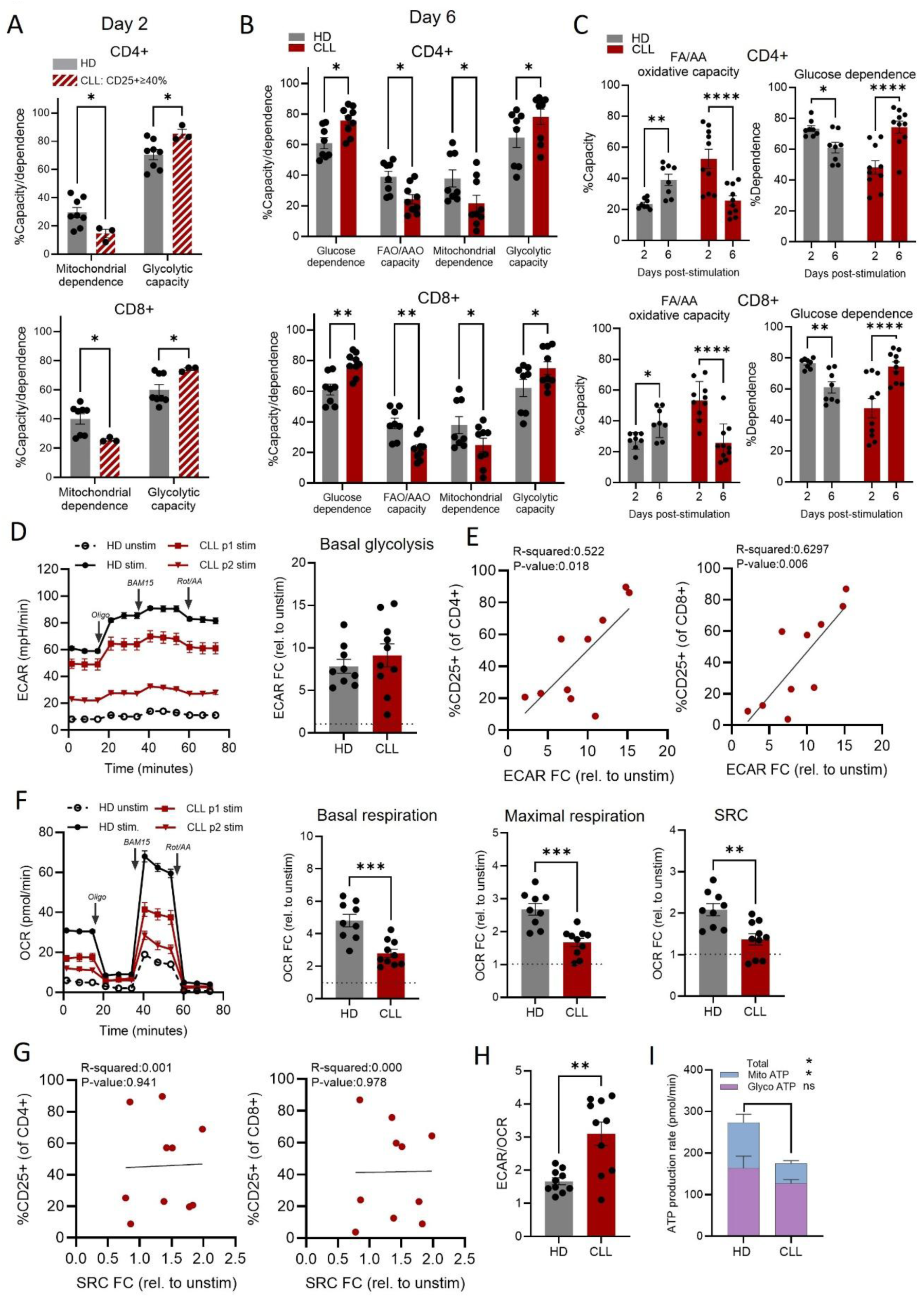
Persistent glycolysis and impaired mitochondrial activity characterize activated CLL T cells. PBMCs from CLL patients or age-matched HDs were stimulated with αCD3/CD28 antibodies and analyzed at day two or day six. (A) Metabolic profile of stimulated T cells at day two. CLL patients were classified into high activation status based on a threshold of 40% CD25⁺ cells in the CD4⁺ compartment. (B) Metabolic profile of stimulated T cells at day six. (C) Changes in the metabolic profile of T cells from day two to day six post-stimulation. (D) Representative extracellular acidification rate (ECAR) profiles of T cells isolated from PBMCs of one HD and two representative CLL patients (P1 and P2). PBMCs were left unstimulated (unstim.) or stimulated for two days prior to Seahorse analysis. Basal glycolysis is presented as fold change (FC) relative to unstim. (E) Correlation analyses between ECAR and CD25⁺ cell percentage in CD4⁺ and CD8⁺ T cells. (F) Representative oxygen consumption rate (OCR) profiles of T cells isolated from PBMCs of one HD and two representative CLL patients (P1 and P2). PBMCs were left unstimulated (unstim.) or stimulated for two days prior to Seahorse analysis. Basal respiration, maximal respiration, and spare respiratory capacity (SRC) are presented as FC relative to unstim. (G) Correlation analyses between SRC and CD25⁺ cell percentage in CD4⁺ and CD8⁺ T cells. (H) Ratio of basal ECAR to basal OCR. (I) Total ATP production rate, partitioned into mitochondrial ATP production rate (mitoATP) and glycolytic ATP production rate (glycoATP) (HD: N = 9, CLL: N = 11). Differences were analyzed using multiple unpaired two-tailed t tests (A–C), unpaired two-tailed t tests (D, F, H), two-way ANOVA (I), or simple linear regression (E, G). A p-value < 0.05 was considered statistically significant. *p < 0.05; **p < 0.01; ***p < 0.001; ****p < 0.0001. Error bars represent mean ± SEM.

As a complementary approach, extracellular flux analysis was performed on T cells isolated from HD or CLL PBMCs after two days of stimulation. Glycolytic activity and mitochondrial oxidative phosphorylation (OXPHOS) were assessed through extracellular acidification rate (ECAR) and oxygen consumption rate (OCR), respectively. HD T cells upregulated glycolysis upon stimulation, whereas CLL T cells showed a heterogeneous glycolytic response that correlated with activation levels (Fig. 2D-E). In contrast, mitochondrial respiration was impaired in CLL T cells, with reduced basal respiration, maximal respiration, and spare respiratory capacity (SRC), irrespective of activation levels (Fig. 2F–G). This was reflected in an increased ECAR/OCR ratio (Fig. 2H) and reduced total ATP production due to diminished mitochondrial ATP generation in patients (Fig. 2I). Together, these findings confirm CENCAT results and demonstrate a persistent glycolytic phenotype with impaired mitochondrial function in CLL T cells.

Effector skewing has been described in the CD8+ compartment of CLL patients^14,15^. Indeed, an enrichment in terminally differentiated effector memory T cells (CD45RA+CD27–, EMRA) was confirmed in the CD8+ compartment of CLL patients included in our CENCAT analyses. Distribution of the CD4+ compartment was not different from HD (Fig. S2D). In these patients, a bias towards glucose metabolism at the expenses of mitochondrial oxidation was observed in both CD4⁺ and CD8⁺ cells (Fig. 2A-C). This argues against subset skewing as the sole explanation for altered metabolic responses in CLL T cells and instead suggests broader, subset-independent metabolic dysregulation.

### Hyporesponsive T cells display reduced glutamine use in the mitochondria

We next examined whether differences in fuel uptake or utilization could explain the low mitochondrial activity identified in T cells from CLL patients. Surface expression of transporters of the two essential fuels for T cell activation, glucose (GLUT1) and glutamine (ASCT2), was measured in T cells stimulated for two days. T cells from CLL patients with poor activation responses (CD25+<40%) showed reduced expression of both GLUT1 and ASCT2, while T cells from patients with preserved activation capacity (CD25+≥40%) had levels comparable to HD (Fig. 3A). We then assessed fuel utilization using 13C-labeled glucose and glutamine. In stimulated T cells from both HD and CLL, glucose was preferentially metabolized via glycolysis, while glutamine was the main fuel of the tricarboxylic acid (TCA) cycle. Incorporation of glutamine in the TCA cycle increased with T cell activation in CLL. As an example, 13C labelling from glutamine in aspartate was significantly decreased in T cells from CLL patients with low activation levels, while it was not different from HD in T cells from patients with preserved activation capacity (Fig. 3B).

**Figure 3.**
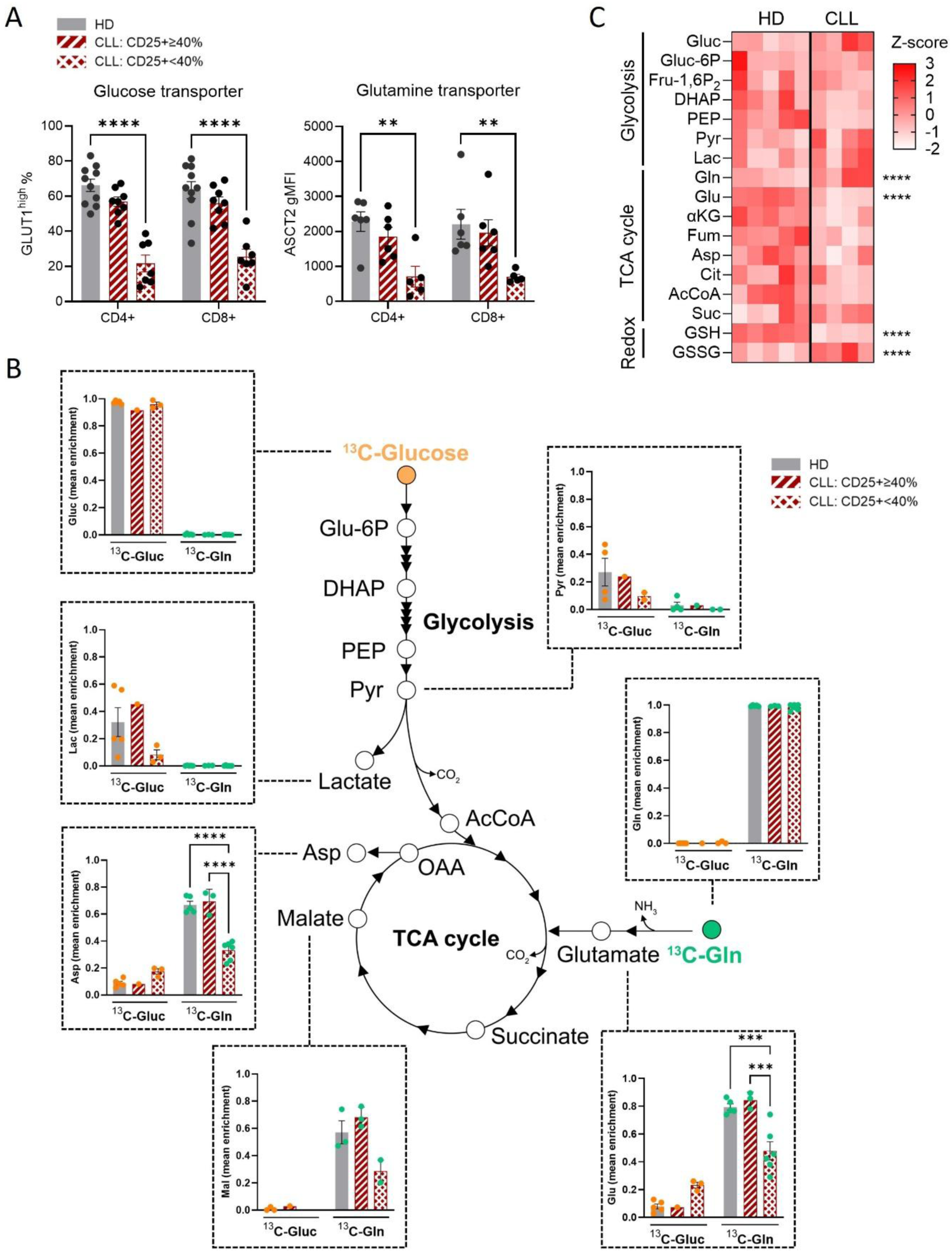
Hyporesponsive T cells display reduced glutamine use in the mitochondria. PBMCs from CLL patients or age-matched HDs were stimulated with αCD3/CD28 antibodies for two days. (A) Cell surface expression levels of the glutamine transporter ASCT2 and glucose transporter GLUT1 in T cells. CLL patients were classified into high and low activation status based on a threshold of 40% CD25⁺ cells in the CD4⁺ compartment. (B) LC–MS/MS analysis of glycolytic and TCA cycle intermediates in T cells isolated from HDs or CLL PBMCs and cultured in medium containing fully labeled ¹³C-glucose (orange) or ¹³C-glutamine (green). Mean enrichment indicates the fraction of ¹³C-labeled molecules of a given metabolite relative to the total amount of that metabolite within a sample (HD: N = 5, CLL: N = 9). Sample size differs per metabolite due to limitations in annotation of certain MS peaks across experiments. (C) Metabolomics analysis of T cells isolated from HDs (N = 5) or CLL PBMCs (N = 4). Heatmap showing the abundance of metabolites from glycolysis, the TCA cycle, and stress response pathways. Abundance is shown as Z-score. Differences were analyzed using two-way ANOVA. A p-value < 0.05 was considered statistically significant. *p < 0.05; **p < 0.01; ***p < 0.001; ****p < 0.0001. Error bars represent mean ± SEM.

Consistent with 13C isotope labelling results, metabolomics showed differences between T cells from HD and hyporesponsive CLL T cells (Fig. S3A, S3B), with a trend towards decreased metabolite abundance across several pathways in CLL (Fig. S3C). Interindividual variability in the abundance of glycolytic intermediates was found in HD and CLL, with no significant differences between the two groups. According to 13C-glutamine isotope labelling, a trend towards decreased levels of all TCA cycle intermediates was seen in CLL T cells (Fig. 3C). Glutamine was significantly higher in CLL T cells compared to HD while glutamate was significantly lower, compatible with decreased glutaminolysis in T cells from hyporesponsive CLL patients. Additionally, reduced glutathione (GSH) was significantly decreased while oxidized glutathione (GSSG) was significantly increased in T cells from CLL, suggesting an stress response occurring in these cells.

In order to assess the abundance of TCA cycle intermediates in T cells from CLL patients with preserved activation levels, intracellular malate concentration was measured in a targeted assay. Malate abundance correlated with activation levels (CD25 expression) in CLL T cells (Fig. S3D) and was not different between HD and CLL (Fig. S3E), further supporting the notion that TCA cycle fueling is not limiting in T cells with high activation levels.

Overall, these results indicate that reduced glutamine uptake and metabolization limit OXPHOS in hyporesponsive CLL T cells, but not in T cells with high activation, which still display low OXPHOS. Thus, subsequent analyses focused exclusively on activation-proficient CLL patients (CD25+ ≥ 40%) in order to investigate the basis of low OXPHOS in highly activated T cells.

### Impaired OXPHOS in activated CLL T cells is associated with altered mitochondrial architecture

To specifically assess oxygen consumption in the electron transport chain (ETC) independent of fuel availability and cytoplasmic metabolism, we performed extracellular flux analysis in conditions of plasma membrane permeabilization (PMP) with saturating mitochondrial substrates and ADP. CLL T cells showed significantly reduced respiration compared to HD T cells also in these conditions (Fig. 4A), pointing to alterations in the activity of the ETC. Maximum uncoupled respiration following FCCP injection was comparable to basal respiration, indicating that OCR is not constrained by ATP synthase (complex V) activity in CLL T cells, but may reflect a general decrease in the efficiency of the ETC.

**Figure 4.**
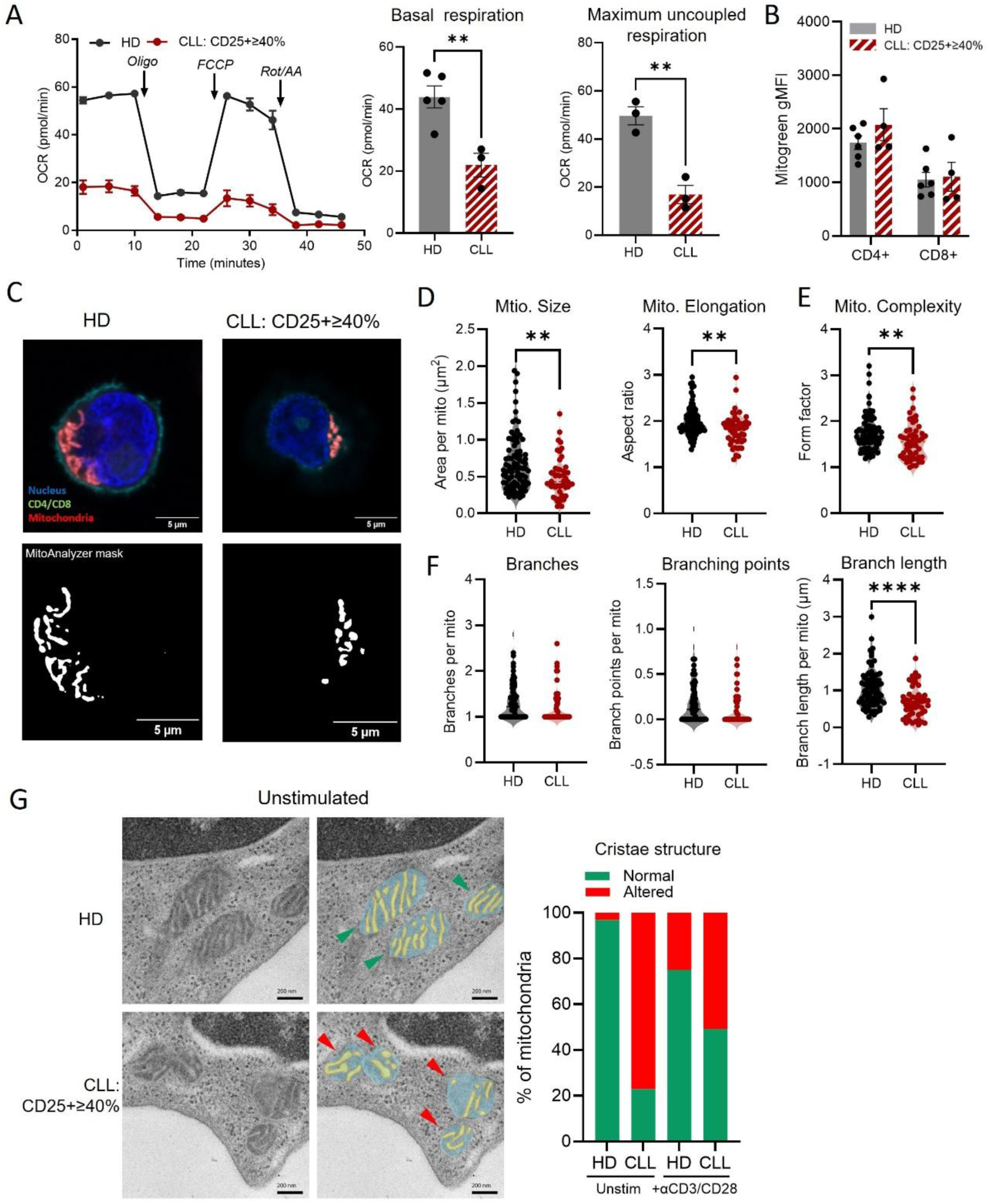
Impaired OXPHOS in activated CLL T cells is associated with altered mitochondrial architecture. PBMCs from CLL patients or age-matched HDs were stimulated with αCD3/CD28 antibodies for two days. (A) Representative oxygen consumption rate (OCR) profile. OCR was measured in permeabilized T cells isolated from HD or CLL PBMCs after stimulation. Mitochondrial substrates and ADP were provided in excess to drive phosphorylating respiration, and FCCP was injected to determine maximal uncoupled respiration. (B) Mitochondrial mass measured by MitoTracker Green. (C) Confocal imaging of mitochondria in T cells isolated from HD and CLL PBMCs after stimulation. Morphological analysis was performed using the Mitochondria Analyzer (Fiji). (HD: N = 2, CLL: N = 2). (D–F) Quantification of mitochondrial structural parameters; each dot represents a single cell: (D) mitochondrial size (mito. size) and elongation, (E) mitochondrial complexity, (F) number of branches, branching points, and branch length per mitochondrion. (G) Transmission electron microscopy of T cells isolated from HD and PBMCs after stimulation. (HD: N = 1, CLL: N = 2=1). Blue: mitochondrial matrix, yellow: cristae, red: inner mitochondrial membrane. Percentage of mitochondria classified based on cristae morphology: normal (parallel, thin cristae homogeneously distributed) or altered (disorganized and/or swollen cristae with heterogeneous distribution). Scoring was performed blindly by three independent evaluators. Differences were analyzed using unpaired two-tailed t tests (A, D–F) and two-way ANOVA (B). A p-value < 0.05 was considered statistically significant. *p < 0.05; **p < 0.01; ***p < 0.001; ****p < 0.0001. Error bars represent mean ± SEM.

One explanation for reduced mitochondrial activity can be reduced mitochondrial mass. However, no differences were observed between HD and CLL T cells (Fig. 4B). Instead, confocal imaging revealed differences in mitochondrial architecture upon stimulation (Fig. 4C). Quantitative analysis showed that mitochondria in CLL T cells were significantly smaller and exhibited reduced elongation, as measured by aspect ratio, compared to HD T cells (Fig. 4D; Fig. S4A). Mitochondrial structural complexity, assessed by form factor, was decreased in CLL T cells, reflecting less connectivity (Fig. 4E;Fig. S4A). Although the number of mitochondrial branches and branching points was similar in the two groups, branch length was markedly reduced in CLL T cells, suggesting impaired network extension (Fig. 4F). At baseline, none of the mitochondrial parameters were significantly different between the two groups, only mitochondrial elongation showed a trend toward reduction in T cells from CLL patients (Fig. S4B-C).

Given the tight link between mitochondrial structure, cristae organization and ETC efficiency^24^, we examined cristae ultrastructure by electron microscopy. In HD T cells, mitochondrial cristae appeared thin and arranged in parallel stacks (Fig. 4G), consistent with efficient ETC assembly in line with the high OXPHOS observed by extracellular flux analyses. In contrast, CLL T cells had more swollen and disorganized cristae (Fig. 4G), which have been previously associated with impaired ETC organization and reduced OXPHOS in effector T cells incapable of mitochondrial fusion^24^. Cristae abnormalities were already present in CLL T cells before stimulation and persisted following CD3/CD28 engagement, indicating a pre-existing impairment in mitochondrial ultrastructure that persists upon stimulation. Nevertheless, mitochondrial with normal cristae were more frequently identified after stimulation that at baseline in CLL. This, together with evidence of increased mitochondrial biogenesis (Fig. S4D) suggests that T cells from CLL patients retain the capacity to produce new functional mitochondria upon stimulation, while a substantial fraction of altered mitochondria remains.

### mTORC1 activity sustains glycolytic bias and hyperactivation in CLL T cells

At the core of T-cell metabolic reprogramming, mTORC1 coordinates the early glycolytic shift in effector cells and induces mitochondrial biogenesis, protein synthesis and proliferation^25–28^. We found that CLL T cells displayed increased expression of mTORC1-associated genes at early time points after stimulation, as revealed by interrogation of the Hallmark MTORC1 SIGNALING gene set in RNA-seq analysis of T cells isolated from HD or CLL PBMCs after two days of stimulation (Fig. 5A; Fig. S5A). Consistently, c-Myc transcriptional targets were upregulated, including several genes involved in the regulation of protein translation and glycolysis (Fig. S5A). To measure the activity of mTORC1 at the protein level, we assessed the phosphorylation of ribosomal protein S6 (p-S6). p-S6 levels were not different between HD- and CLL-derived T cells after two days of stimulation, however at day six T cells from CLL PBMCs showed significantly higher p-S6 (Fig. 5B). In HD, instead, a clear downregulation of p-S6 was observed from day two onto day six. In line with the described role of mTORC1 in translation initiation^29^, protein synthesis rate was significantly increased in CLL T cells (Fig. 5C). p-S6 levels progressively increased from low- to high-proliferating CLL T cells (Fig. 5D), consistent with the role of mTORC1 supporting proliferation.

**Figure 5.**
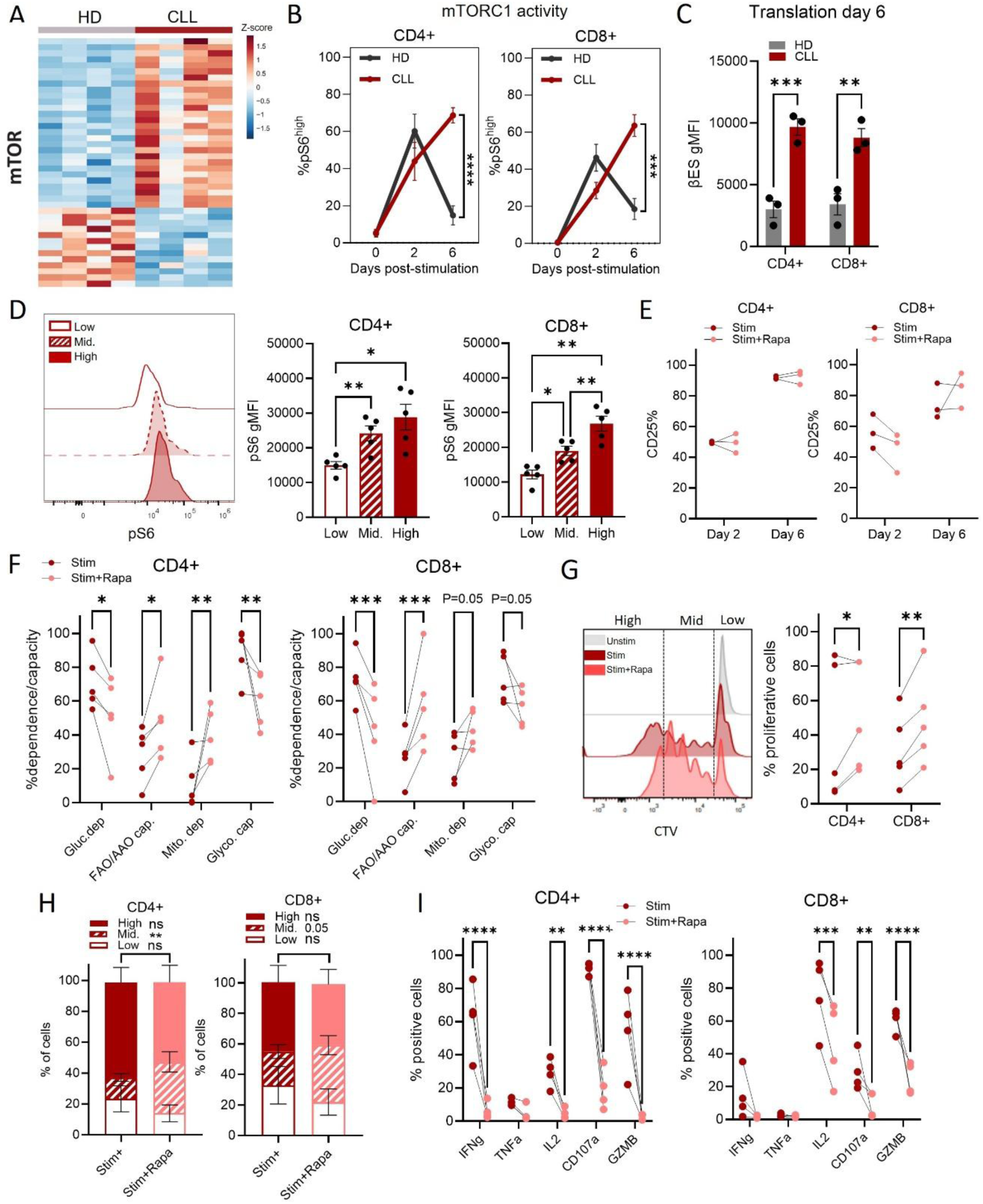
mTORC1 activity sustains glycolytic bias and hyperactivation in CLL T cells. PBMCs from CLL patients or age-matched HDs were stimulated with αCD3/CD28 antibodies for two days (A) or six days (B–I). (A) Heatmap depicting differentially expressed genes (padj < 0.05) associated with the mTORC1 pathway (Hallmark gene set MTORC1 SIGNALING) in T cells isolated from HD and CLL PBMCs. (B) mTORC1 activity measured as the percentage of cells expressing high p-S6 levels at the indicated time points (HD: N = 7, CLL: N = 7). (C) Protein translation measured by βES incorporation into newly synthesized proteins. (D) p-S6 levels in low-, intermediate-, and high-proliferating T cells from CLL patients. (E) Percentage of CD25⁺ T cells in CLL T cells at day two and six post-stimulation in the presence or absence of rapamycin. (F) Metabolic profile of CLL T cells in response to stimulation in the presence or absence of rapamycin. (G) Representative proliferation profile of CD4⁺ T cells measured by CellTrace Violet (CTV) and example of the gating strategy. Quantification of the percentage of cells that entered division (% proliferating cells) and distribution of cells undergoing low (0–1 divisions), intermediate (2–4 divisions), or high (>5 divisions) proliferation based on the indicated gating strategy. (H) Distribution of cells undergoing low, intermediate, or high proliferation (N = 5). (I) Percentage of cytokine-positive and degranulating (CD107a⁺) T cells upon stimulation in the presence or absence of rapamycin, determined by intracellular cytokine staining and surface CD107a staining. Differences were analyzed using one-way ANOVA (D), two-way ANOVA (B), or multiple paired two-tailed t tests (E,C, F–I). A p-value < 0.05 was considered statistically significant. *p < 0.05; **p < 0.01; ***p < 0.001; ****p < 0.0001. Error bars represent mean ± SEM.

To determine whether sustained mTORC1 activity contributes to the observed glycolytic and hyperactivated phenotype identified in CLL T cells, we stimulated CLL PBMCs in the presence of the mTORC1 inhibitor rapamycin. Rapamycin effectively reduced p-S6 levels and global protein translation (Fig. S5B) without impairing the ability of T cells to become activated (Fig. 5E). According to our hypothesis, rapamycin increased mitochondrial dependence and FA/AAO capacity in CLL T cells while reducing glucose dependence and glycolytic capacity (Fig. 5F), indicating that mTORC1 sustains the glycolytic phenotype of CLL T cells. Furthermore, mTORC1 inhibition significantly increased the amount of proliferative cells (Fig. 5G), and led to a more balanced distribution of proliferation cycles (Fig. 5H). In parallel, rapamycin treatment significantly decreased cytokine production, degranulation (CD107a) and expression of GZMB in both CD4+ and CD8+ T cell compartments (Fig. 5I). Together, these findings support a model in which sustained mTORC1 activity contributes to the persistent glycolytic bias and hyperactivated proliferative phenotype of CLL T cells.

### Exogenous glutamine uptake sustains non-lysosomal mTORC1 activity in T cells

mTORC1 activation is classically a two-step process, integrating PI3K–AKT signaling with amino acid sensing at the lysosome (Fig. 6A). In healthy T cells, PI3K–AKT activation is boosted by CD28 co-stimulation^4^. Similarly, in CLL T cells, phosphorylation of S6 only occurs with combined anti-CD3/CD28 stimulation, and not with CD3 alone (Fig. S6A). AKT inhibition at the onset of stimulation reduced mTORC1 activity (Fig. S6B), confirming that PI3K–AKT signaling drives the initial mTORC1 activation downstream anti-CD3/CD28 stimulation. However, PI3K/AKT inhibition during later stage of activation did not reduce p-S6 levels (Fig. 6B), showing that PI3K–AKT signaling initiates mTORC1 activation but other factors contribute to its sustained activation in CLL T cells. In line with this, no difference in phosphorylation levels of AKT between HD and CLL T cells was observed at late stage (Fig. 6C), despite persistently high p-S6 (Fig. 5B).

**Figure 6.**
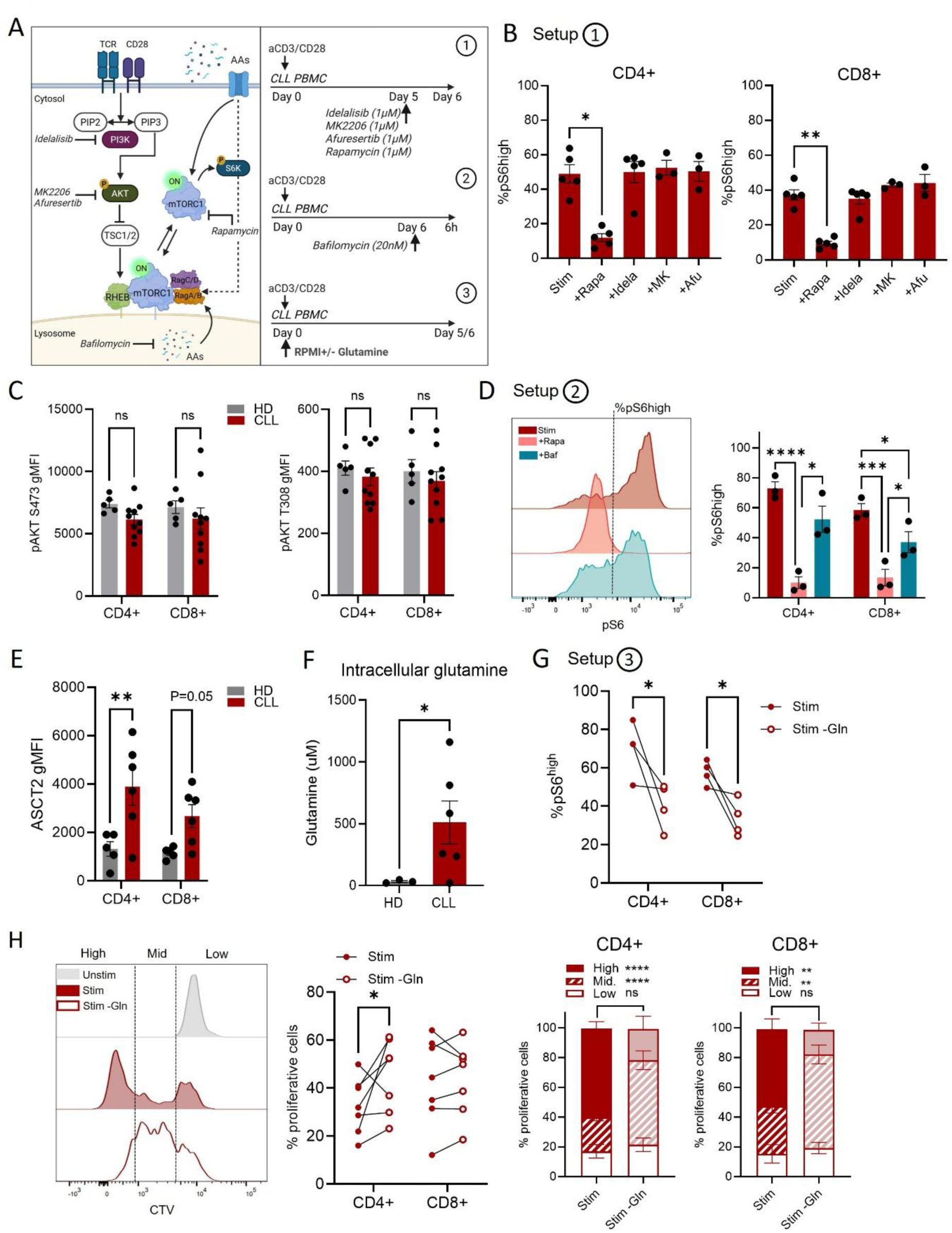
Exogenous glutamine uptake sustains non-lysosomal mTORC1 activity in T cells. PBMCs from CLL patients or age-matched HDs were stimulated with αCD3/CD28 antibodies for six days. (A) Schematic overview of mTORC1 activation (left) and experimental setup using inhibitors and glutamine deprivation (right). (B) mTORC1 activity in CLL T cells measured as the percentage of cells expressing high p-S6 levels after 24-hour treatment on day five with rapamycin (+rapa, 1 µM), idelalisib (+idela, 1 µM), MK2206 (+MK, 1 µM), or afuresertib (+Afu, 1 µM). (C) Phosphorylation of AKT (S473 and T308) in HD and CLL T cells. (D) mTORC1 activity in CLL T cells measured as the percentage of cells expressing high p-S6 levels after six-hour treatment on day six with rapamycin (+rapa, 1 µM) or bafilomycin A1 (+Baf, 20 nM). Representative p-S6 histograms are shown. (E) Surface expression levels of ASCT2 in HD and CLL T cells. (F) Intracellular glutamine levels (µM) in sorted CD4⁺ and CD8⁺ T cells from stimulated HD or CLL PBMCs, lysed at 666 cells/µL. (G) mTORC1 activity in CLL T cells measured as the percentage of cells expressing high p-S6 levels after culture in complete medium or glutamine-free medium (−Gln). (H) Quantification of the percentage of cells that entered division (% proliferating cells) and distribution of cells undergoing low (0–1 divisions), intermediate (2–4 divisions), or high (>5 divisions) proliferation based on the indicated gating strategy (N = 7). Differences were analyzed using one-way ANOVA (B), two-way ANOVA (D, H), multiple unpaired two-tailed t tests (C, E, F), or paired two-tailed t tests (G, H). A p-value < 0.05 was considered statistically significant. *p < 0.05; **p < 0.01; ***p < 0.001; ****p < 0.0001. Error bars represent mean ± SEM.

mTORC1 activation requires amino acids, which can be derived from lysosomal proteolysis or exogenous uptake^30^ (Fig. 6A). Early lysosomal inhibition with Bafilomycin A1 completely blocked p-S6 induction, supporting a requirement for lysosomal function during initial mTORC1 activation (Fig. S6C). In contrast, addition of Bafilomycin A1 during the last 24h of stimulation only partially reduced p-S6 levels in CLL T cells (Fig. 6D). This suggests that sustained p-S6 accumulation in CLL T cells is mostly independent of lysosomal activity. We therefore explored whether extracellular amino acid uptake could contribute to sustained mTORC1 signaling. CLL T cells showed increased surface expression of the glutamine transporter ASCT2 at day six (Fig. 6E), consistent with enhanced capacity for extracellular glutamine uptake. Among the amino acids sensed by mTORC1, only glutamine levels were significantly elevated in CLL T cells at day two (Fig. S6D) and remained elevated at day six (Fig. 6F). Together, these observations support a model in which extracellular glutamine uptake contributes to sustained mTORC1 activity in CLL T cells.

To further investigate the contribution of extracellular glutamine availability to sustained mTORC1 activity, we depleted glutamine from the culture medium during stimulation of CLL PBMCs. Glutamine deprivation significantly reduced p-S6 in T cells from CLL patients (Fig. 6G) without affecting T-cell activation levels (Fig. S6E). Accordingly, glutamine deprivation normalized the proliferation profile of CLL T cells. Mainly, it increased the percentage proliferative cells in the CD4+ compartment and decreased highly proliferative cells in both CD4+ and CD8+ compartments (Fig. 6H).

Together, these findings support a role for extracellular glutamine uptake in sustaining mTORC1 activity and altered proliferative responses in CLL T cells. These data further support a model of persistent metabolic hyperactivation in dysfunctional CLL T cells.

## Discussion

Our study identifies a hyperactivated state in tumor-associated T cells from CLL patients that contrasts with the conventional view of cancer-induced T-cell dysfunction. Upon stimulation, these cells exhibit excessive division cycles and increased cytokine production, sustained by mTORC1 activity and a persistent glycolytic bias. Concomitantly, OXPHOS is reduced and mitochondrial structure becomes disrupted. We propose a model in which reduced OXPHOS reflects both mitochondrial defects that pre-exist in T cells from CLL patients before TCR engagement, and a failure to engage mitochondrial metabolism upon activation. This builds on and further extends our previous finding that mitochondrial depolarization is already present before stimulation in CLL T cells, even in naïve and central memory compartments^17^. Upon stimulation, sustained mTORC1 activity further reinforces the pre-existent OXPHOS limitation by maintaining high glycolytic activity and limited mitochondrial engagement. The resulting hyperactivated state is likely unsustainable, as reflected by the failed response to repeated stimulation in CLL T cells reported earlier by our group^18^. This is consistent with previous work showing that genetically induced mitochondrial dysfunction in T cells promotes inflammatory programs while limiting T-cell survival^8,31^, and that intact mitochondrial respiration is required for memory T-cell differentiation^24,32,7^. Besides, constitutive activation of the mTORC1 pathway has been reported in individuals carrying the pathogenic m.3243A>G mtDNA mutation, supporting the notion that mitochondrial dysfunction can directly enforce aberrant mTORC1 signaling^33^.

In solid tumors, metabolic interventions that increase the expression of fuel transporters have been proposed to improve T-cell metabolic activity and anti-tumor function of CAR-T cells^34,35^. However, these approaches are designed to overcome hypo-responsive metabolic states and could potentially exacerbate the hyperactivated phenotype observed in CLL T cells, which already display high nutrient uptake. Instead, we find that impaired oxidative metabolism in CLL T cells was associated with abnormalities in mitochondrial structure, including reduced mitochondrial complexity and altered cristae architecture. A decreased GSH/GSSG ratio further supports increased oxidative stress in these cells, consistent with our previous findings^21^. Our observations are in line with studies linking mitochondrial fragmentation and disrupted cristae to glycolytic phenotype, whereas fused mitochondrial networks have been described to support efficient oxidative metabolism and memory development in T cells^24^. In line with this, accumulation of dysfunctional mitochondria in TILs has been shown to impair anti-tumor responses^36^. Cristae abnormalities were detected in CLL T cells prior to stimulation and persisted after activation despite intact mitochondrial biogenesis. The coexistence of newly formed mitochondria with regular structure and a persistent pool of mitochondria with altered architecture might suggest defects in mitochondrial quality control and clearance pathways. Whether this involves impaired mitophagy remains to be investigated.

mTORC1 is a central regulator of T cell fate decisions, integrating environmental signals to shape metabolic programming^37^. Previous studies have shown that mTORC1 promotes anabolic metabolism and remodels mitochondrial function by inhibiting mitophagy and promoting fission^38,39^. Although this program supports early effector function, sustained mTORC1 activity limits metabolic flexibility and compromises long-term T cell responses during chronic infection^40^. Notably, prolonged mTORC1 signaling has been associated with impaired memory formation^41,42^, whereas enhancing mitochondrial quality control by mitophagy promotes the generation of memory T cells^36,43^. Indeed, we find that sustained mTORC1 signaling in CLL T cells is linked with persistent reliance on glycolysis and a hyperactivated state. Additionally, mTORC1 inhibition in our model restores mitochondrial engagement upon anti-CD3/CD28 stimulation, reduces hyperactivation and normalizes proliferation dynamics. This is in line with studies performed in acute infections showing that mTOR inhibition promotes memory formation^44^. More recently, rapamycin has been shown to reduce exhaustion and lead to smaller viral reservoir in mice infected with HIV and treated with CAR-modified hematopoietic stem cells^45^.The signaling vulnerability identified here may underlie the skewing of CLL T cells toward highly differentiated effector phenotypes and the loss of naïve and memory T cell populations observed by us and others^14–16^, and might represent a potential target to restore long-term T cell function not only in CLL.

mTORC1 is classically described as a downstream effector of PI3K–AKT signaling. Constitutive activation of this pathway, as seen in activated PI3Kδ syndrome, drives increased mTORC1 activity, glycolysis, and effector skewing^46,47^. We indeed find that AKT contributes to the initial activation of mTORC1 following TCR stimulation, and that this early phase depends on lysosomal activity, consistent with the canonical model in which amino acid sensing promotes mTORC1 recruitment to the lysosomal surface^48^. At later stages, however, sustained mTORC1 activity appears largely independent of both AKT and lysosomal function. Instead, our data support a role for exogenous glutamine uptake in sustaining mTORC1 activity at later stages of activation. These findings extend recent evidence for non-lysosomal mTORC1 activation^30^ to primary human T cells and suggest that metabolic inputs can contribute to sustained mTORC1 activity beyond canonical upstream signaling.

Collectively, our findings suggest that sustained promotion of effector function and nutrient uptake may not be beneficial across all tumor settings. Thus, metabolic interventions in T-cell therapy should be tailored to the functional state of tumor-associated T cells. In hypo-responsive or exhausted settings, strategies that increase activation may be beneficial, whereas hyperactivated T cells may instead benefit from restraining excessive activation to maintain durability. In line with this concept, recent efforts have focused on limiting T-cell activation and terminal differentiation in CAR-T cells to preserve stem-like states, an approach that improves persistence and anti-tumor efficacy^49–51^.

## Limitations of the study

An important open question is whether the hyperactivated metabolic phenotype described here in T cells from CLL PBMCs is also present in lymph node–resident T cells. In CLL lymph nodes, restricted nutrient and oxygen supplies and local suppressive cues could shape T-cell metabolism differently, as extensively documented in solid malignancies. In this regard, recent work by Llaó-Cid et al. described enrichment of exhausted T cells in CLL lymph nodes as compared with peripheral blood^52^, suggesting that the composition of T cell populations differs across compartments. In addition, we previously showed that lymph node stimuli boost glutamine metabolism in CLL cells^53^, which could change the availability of amino acids for T cells in the vicinity. Whether sustained mTORC1 activity also characterizes lymph node–resident T cells remains to be determined.

Another limitation is related to the intrinsic multicellular nature of our PBMC cultures. Although we show that rapamycin directly inhibits mTORC1 signaling in T cells, its effects on CLL cells within PBMCs may also contribute indirectly, for example by altering CLL-cell proliferation^54^. While direct and indirect effects of rapamycin cannot be fully disentangled in this system, the functional and metabolic changes observed in T cells are consistent with established mTORC1-dependent processes, supporting a model in which sustained mTORC1 signaling contributes to the dysfunctional hyperactivated state observed in T cells from CLL patients.

## Supporting information

Key resources table

## Acknowledgements

The authors are deeply grateful to the patients and healthy donors who generously contributed samples to this study. The authors thank colleagues and collaborators for insightful discussions and critical feedback throughout the project. Specifically, we want to thank Dr. Sander van Kasteren (Leiden University) for help in setting up CENCAT, Prof. Derk Amsen (Sanquin/Amsterdam UMC) and Dr. Maria Themeli (Amsterdam UMC) for useful discussions, Dr. Jeroen Guikema (Amsterdam UMC) for recommendations on studying the AKT/MORC1 pathway and Dr. Fleur Peters (Amsterdam UMC) for help with analysis and interpretation of the RNAseq data.

This work was supported by an ERC Consolidator Grant [864815, BOOTCAMP] and a Lymph and Co Grant [2018-LYCo-008] granted to A.P.K. Additional support was provided by an EHA Research Grant award [RG-202309-04188] and a CCA grant [CCA2023-9-92] granted to H.S.M.

## Author contributions

NBG, HSM, EE and APK designed the experiments; NBG, GC, EC, HSM, KV, AEG, CFJ,YH, BS and MvW performed experiments; FV and KMB provided material for CENCAT and contributed to its setup and troubleshooting; NBG, HSM, EC, GC, KV, AEG, CFJ, YH, NvdW, BS and MvW analyzed data; APK and MDL provided primary material; NBG and HSM wrote the original draft; NBG, HSM, EE and APK reviewed and edited the manuscript; HSM, EE and APK supervised the project; APK and HSM conceptualized the research, acquired funding and managed the project.

## Resource availability

### Lead contact

Further information and requests for resources and reagents should be directed to and will be fulfilled by the lead contacts, Dr. Helga Simon-Molas and Prof. Arnon P Kater.

### Materials availability

This study did not generate new, unique reagents.

### Data and code availability

- Metabolomics and 13C isotope labelling data will be made available in the open repository MetaboLights (EMBL-EBI).
- The RNA-seq data generated in this study have been deposited in the European Genome-phenome Archive (EGA) under accession number EGAD50000001368.
- This paper does not report original code.
- Any additional information required to reanalyze the data reported in this work paper is available from the lead contacts upon request.

## Supplemental figures

**Figure S1.**
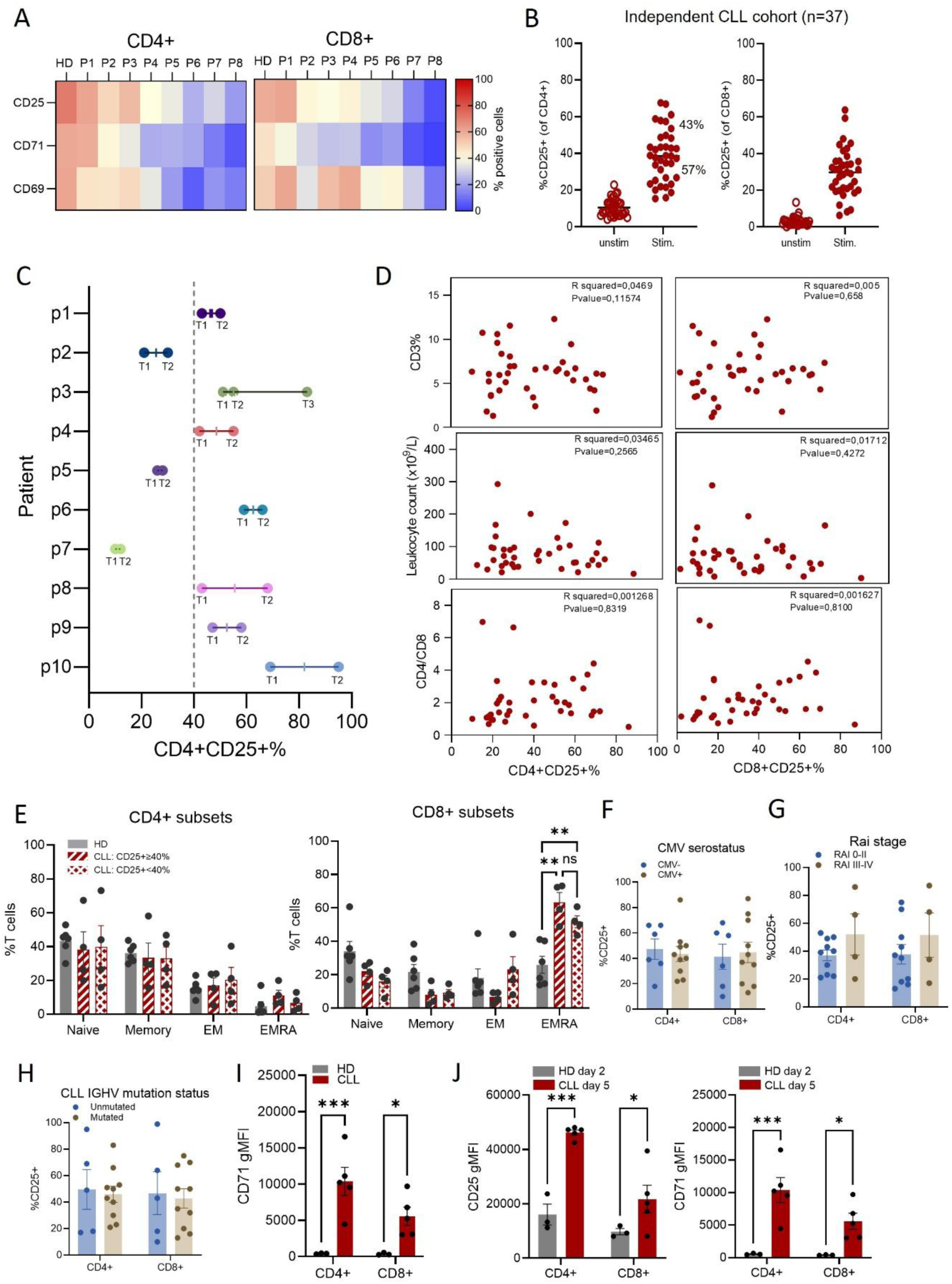
PBMCs from CLL patients or age-matched HDs were analyzed at baseline (day zero; D–H) or after stimulation with αCD3/CD28 for two (A–H) or five days (I). (A) Percentage of CD25⁺, CD71⁺, and CD69⁺ T cells in HDs (mean of N = 3) and individual CLL patients (P1–P8). (B) Percentage of CD25⁺ T cells in an independent cohort of untreated CLL patients (N = 37). (C) Longitudinal analysis of the percentage of CD25⁺ T cells in PBMCs from the same patients (p1-p10) at different time points during disease progression (T1-T3). (D) Correlation analyses of baseline leukocyte count, CD3⁺ T-cell percentage, and CD4/CD8 ratio with the percentage of CD25⁺ cells after stimulation. (E) Baseline T-cell subset distribution at baseline, defined as naïve (CD45RA⁺CD27⁺), memory (CD45RA⁻CD27⁺), effector memory (EM; CD45RA⁻CD27⁻), and EMRA (CD45RA⁺CD27⁻). CLL patients were classified into high and low activation status based on a threshold of 40% CD25⁺ cells in the CD4⁺ compartment. (F) CMV serostatus (CMV⁻/⁺). CLL patients were classified into high and low activation status based on a threshold of 40% CD25⁺ cells in the CD4⁺ compartment. (G) IGHV mutation status (unmutated/mutated). CLL patients were classified into high and low activation status based on a threshold of 40% CD25⁺ cells in the CD4⁺ compartment. (H) Rai stage (0–II vs III–IV). CLL patients were classified into high and low activation status based on a threshold of 40% CD25⁺ cells in the CD4⁺ compartment. (I) CD71 expression levels. (J) CD25 and CD71 expression levels in HD T cells at day two and CLL T cells at day 5 post-stimulation. Differences were analyzed using two-way ANOVA (E–J), multiple unpaired two-tailed t tests (I), or simple linear regression (D). A p-value < 0.05 was considered statistically significant. *p < 0.05; **p < 0.01; ***p < 0.001; ****p < 0.0001. Error bars represent mean ± SEM.

**Figure S2.**
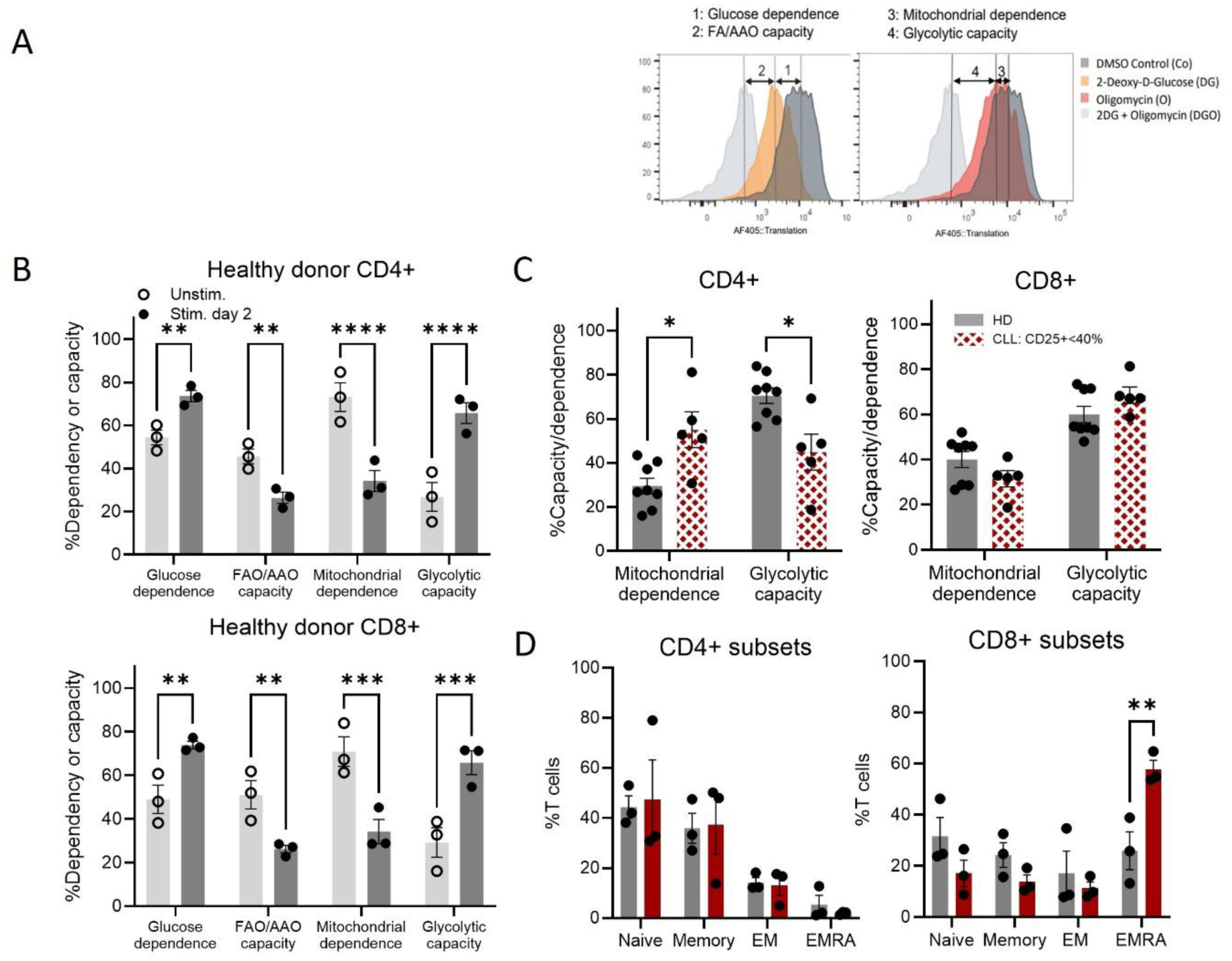
PBMCs from CLL patients or age-matched HDs were stimulated with αCD3/CD28 antibodies and analyzed two days post-stimulation by CENCAT. (A) Schematic overview of the CENCAT approach, illustrating how distinct metabolic pathways contribute to ATP production to support protein translation. Incorporation of βES into newly synthesized proteins is used to quantify metabolic dependencies and capacities under control and inhibitor-treated conditions. Glucose and mitochondrial dependence were calculated as the difference between control and 2-DG- or oligomycin-treated samples, respectively. Fatty acid/amino acid oxidation (FAO/AAO) and glycolytic capacities were derived by subtracting these dependencies from total translation activity. (B) Metabolic profile of HD T cells under unstimulated (unstim.) or stimulated conditions. (C) Metabolic profile of HD and CLL T cells with low activation (CD25 < 40%). (D) T-cell subset distribution at baseline, defined as naïve (CD45RA⁺CD27⁺), memory (CD45RA⁻CD27⁺), effector memory (EM; CD45RA⁻CD27⁻), and EMRA (CD45RA⁺CD27⁻). Measured in patients included in CENCAT analysis in Fig. 2A-C. Differences were analyzed using multiple unpaired two-tailed t tests. A p-value < 0.05 was considered statistically significant. *p < 0.05; **p < 0.01; ***p < 0.001; ****p < 0.0001. Error bars represent mean ± SEM.

**Figure S3.**
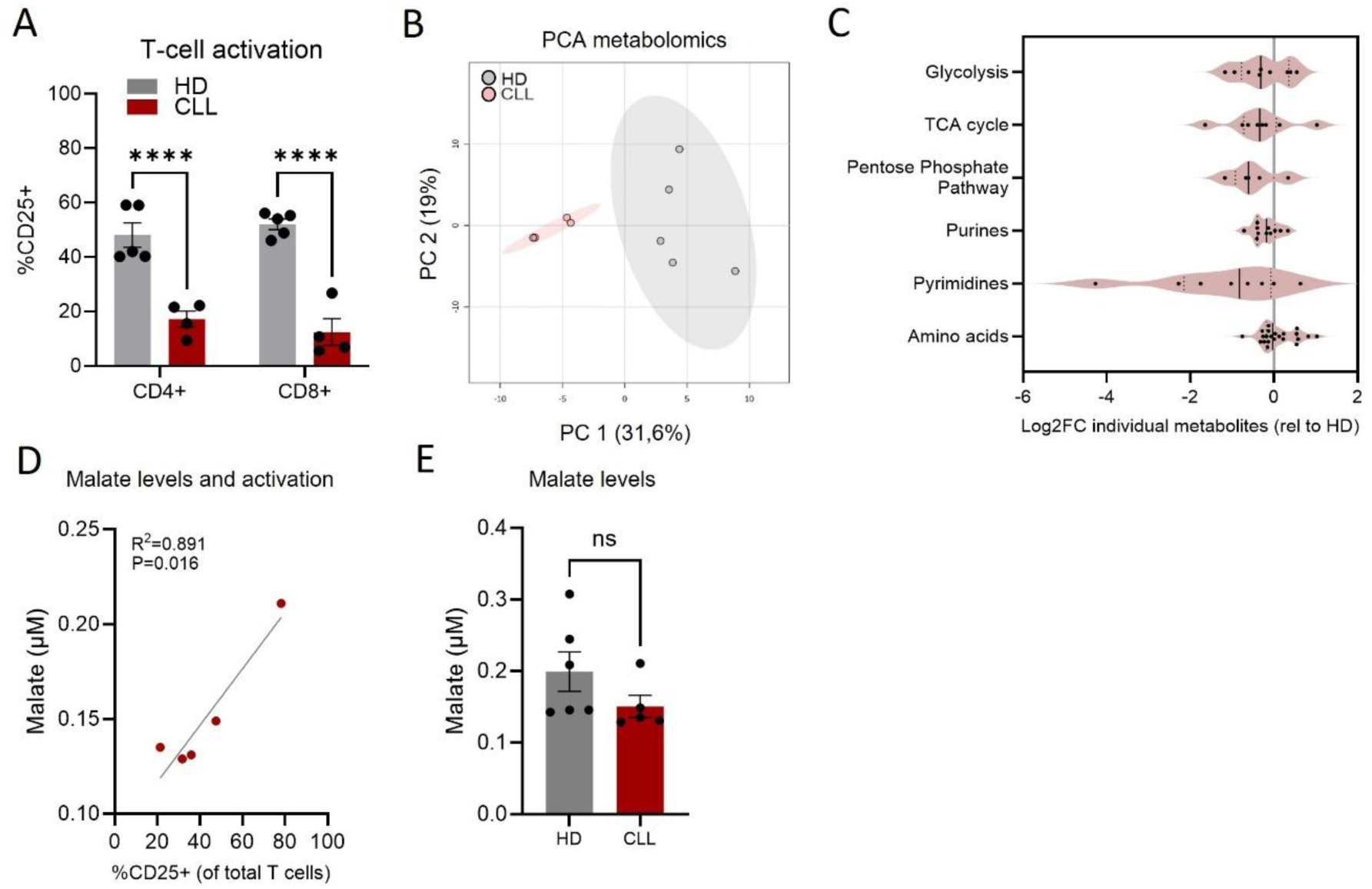
PBMCs from CLL patients or age-matched HDs were stimulated with αCD3/CD28 antibodies for two days. (A) Percentage of CD25⁺ T cells in the samples used for metabolomics analysis. (B) Principal component analysis (PCA) of metabolomics data from T cells isolated from PBMCs after two days of stimulation. (C) Metabolomics analysis of T cells isolated from HDs or CLL PBMCs. Metabolites were grouped by pathway and represented in landscape plots as the mean fold change in metabolite abundance in CLL T cells (N = 4) compared to HD T cells (N = 5). Fold-change abundance of each metabolite in T cells from CLL relative to HD was calculated and represented in violin plots with metabolites grouped by pathway. (D) Intracellular malate levels (µM) measured in T cells isolated from PBMCs, lysed at 666 cells/µL. Correlation analysis of intracellular malate levels against the percentage of CD25⁺ T cells. intracellular malate measured in T cells isolated from PBMCs, lysed at 666 cells/µL. (E)) Intracellular malate levels (µM) measured in T cells isolated from PBMCs, lysed at 666 cells/µL. Differences were analyzed using multiple unpaired two-tailed t tests (A), one-way ANOVA (E), or simple linear regression (D). A p-value < 0.05 was considered statistically significant. *p < 0.05; **p < 0.01; ***p < 0.001; ****p < 0.0001. Error bars represent mean ± SEM.

**Figure S4.**
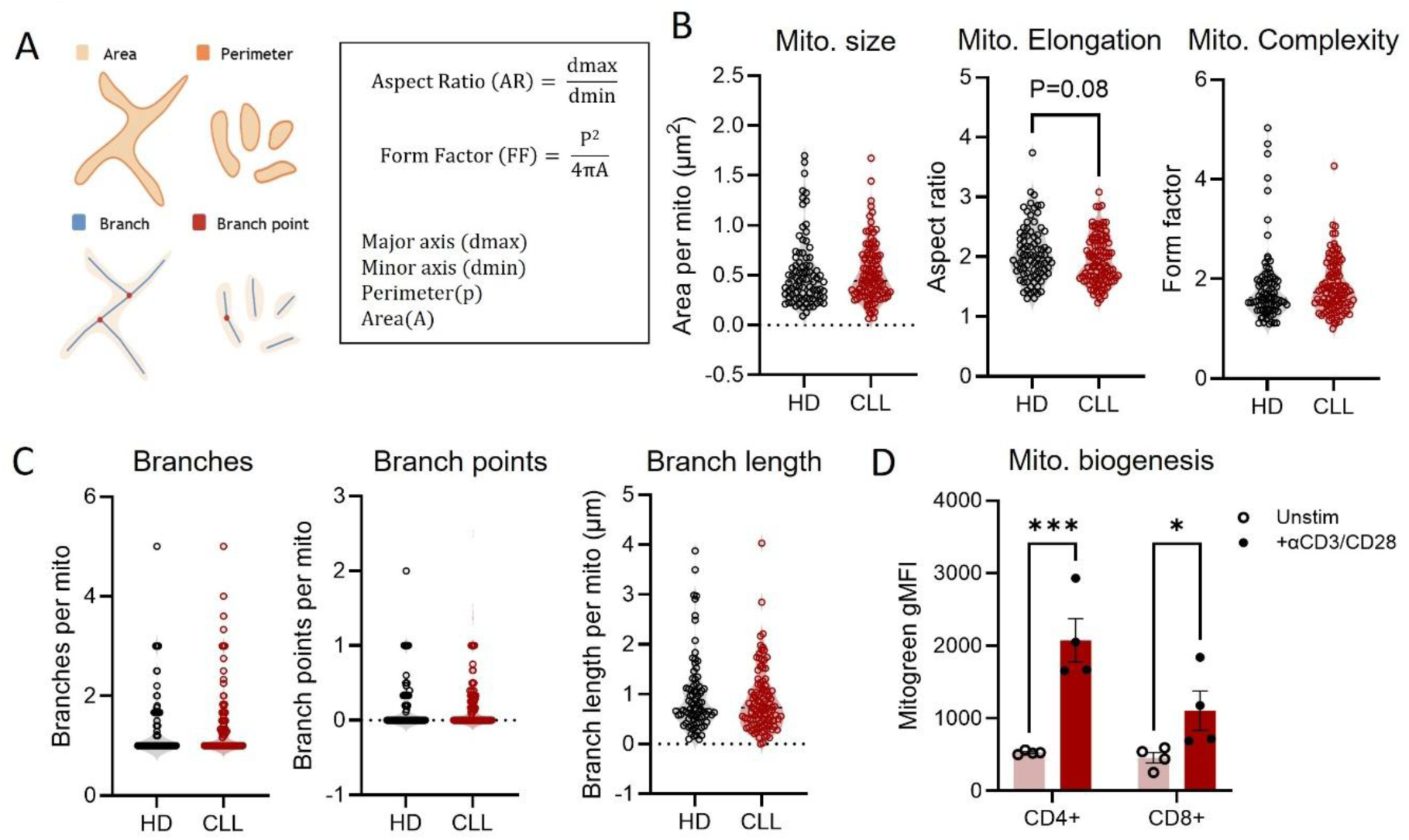
T cells were isolated from PBMCs of HDs or CLL patients after two days without stimulation. (A) Schematic overview of parameters determined using the Mitochondria Analyzer (Fiji), including calculations for aspect ratio and form factor. (B) Quantification of mitochondrial size (mito. size), elongation, and complexity. (C) Quantification of the number of branches, branching points, and branch length per mitochondrion. (D) Mitochondrial mass, measured by MitoGreen, in unstimulated and stimulated CLL T cells at day two post-stimulation. Differences were analyzed using unpaired two-tailed t tests (B-C) and two-way ANOVA (D). A p-value < 0.05 was considered statistically significant. *p < 0.05; **p < 0.01; ***p < 0.001; ****p < 0.0001. Error bars represent mean ± SEM.

**Figure S5.**
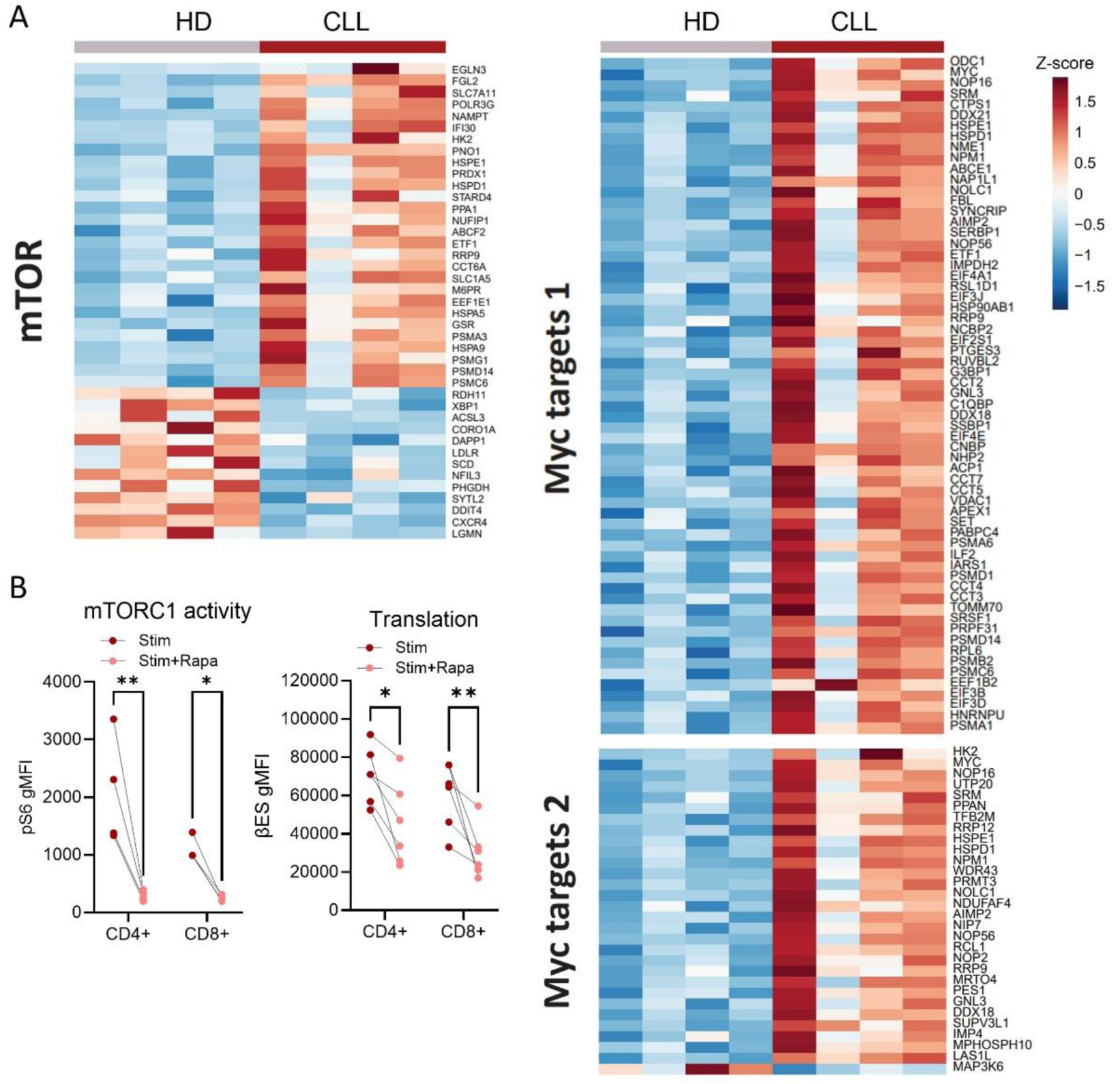
PBMCs from CLL patients or age-matched HDs were stimulated with αCD3/CD28 antibodies for two or six days. (A) Heatmap depicting differentially expressed genes (padj < 0.05) associated with the mTORC1 pathway (Hallmark gene set MTORC1 SIGNALING) and MYC target gene sets (Hallmark gene sets MYC TARGETS V1, and MYC TARGETS V2) in T cells isolated from HD and CLL PBMCs stimulated for two days. (B) mTORC1 activity in CLL T cells measured as the percentage of cells expressing high p-S6 levels and protein translation in CLL T cells at day six post-stimulation in the presence or absence of rapamycin. Differences were analyzed using paired two-tailed t tests (B, C). A p-value < 0.05 was considered statistically significant. *p < 0.05; **p < 0.01; ***p < 0.001; ****p < 0.0001. Error bars represent mean ± SEM.

**Figure S6.**
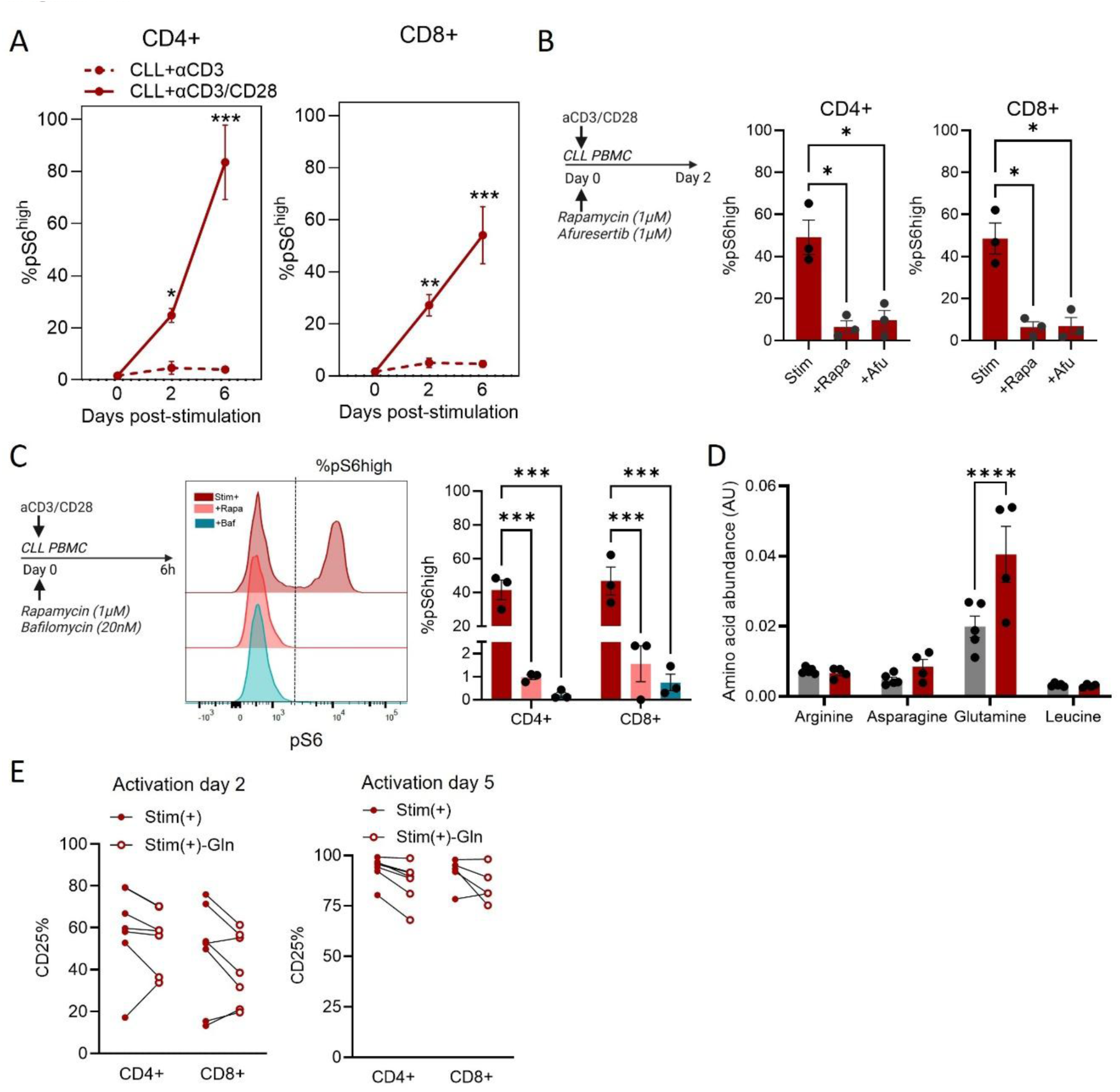
(A) mTORC1 activity measured as the percentage of cells expressing high p-S6 levels in response to stimulation with αCD3 alone or αCD3/CD28 antibodies after two or six days. (B) Percentage of cells expressing high p-S6 levels at day two post-stimulation with αCD3/CD28 antibodies in the presence of rapamycin (+rapa, 1 µM) or afuresertib (+Afu, 1 µM). (C) Schematic overview of the experimental setup in which cells were stimulated in the presence of rapamycin (+rapa, 1 µM) or bafilomycin A1 (+Baf, 20 nM) for six hours. mTORC1 activity in CLL T cells measured as the percentage of cells expressing high p-S6 levels after treatment. (D) Abundance of amino acids (arbitrary units) in T cells from HDs after two days of stimulation. (E) Percentage of CD25⁺ T cells in CLL T cells measured at day two and day five post-stimulation in the presence or absence of glutamine (−Gln). Differences were analyzed using two-way ANOVA (A), one-way ANOVA (B, C), multiple unpaired two-tailed t tests (D), or paired two-tailed t tests (E). A p-value < 0.05 was considered statistically significant. *p < 0.05; **p < 0.01; ***p < 0.001; ****p < 0.0001. Error bars represent mean ± SEM.

## STAR Methods

### Key resources table

See Excel file Key resources table

### Experimental model and study participant details

#### Patients

Peripheral blood samples from patients with chronic lymphocytic leukemia (CLL) were obtained through the B-cell Malignancies Biobank of Amsterdam UMC (Amsterdam, the Netherlands). Sample collection and use were approved by the Medical Ethics Committee of Amsterdam UMC under protocol number 2013-159. A total of n=53 patients were included (Table S1). The median age at sampling was 71 years (range, 51-91), 51% were male and 49% female. Additional clinical characteristics, including Rai stage, IgHV mutation status, Leukocyte count, %CD5+CD19+, %CD3+, CMV status and treatment status are summarized in Table S1. An independent cohort of untreated CLL patients from the University Hospital of Cologne was included with a total of n=37 patients (Table S2). The median age at sampling was 63 year (range, 46-83), 76% were male and 24% female. All participants provided written informed consent in accordance with the Declaration of Helsinki.

#### Healthy donors

Peripheral blood samples from aged-matched (>60 years old) healthy donors (HD) were obtained from Sanquin Blood Supply Foundation (Amsterdam, the Netherlands) according to institutional guidelines and donor informed consent procedures. A total of n = 42 healthy donors were included. The median age was 65 years (range, 60–75), and 55% were male / 45% female. Details can be found in Table S3.

### Method details

#### Cell culture

Peripheral blood mononuclear cells (PBMCs) were isolated from buffy coats or peripheral blood samples obtained from patients CLL and HD by density gradient centrifugation using Ficoll-Paque™ PREMIUM (VWR). Isolated PBMCs were cryopreserved in fetal bovine serum (Takara) containing 20% DMSO and stored in liquid nitrogen until use.

For experiments, cryopreserved PBMCs were rapidly thawed at 37°C, washed and directly cultured as indicated. Cells were seeded at a density of 3 × 10^6 cells/mL in RPMI 1640 medium (Gibco) containing 2 mM L-glutamine, supplemented with 10% fetal bovine serum (Takara), 1% penicillin-streptomycin (Corning) and 46 µM β-mercaptoethanol and maintained at 37°C in a humidified incubator with 5% CO₂ for two–six days. T cell activation was induced using soluble anti-CD3 monoclonal antibody (91 ng/mL; clone 1XE; Sanquin) together with anti-CD28 monoclonal antibody (3 µg/mL; clone 15E8; Sanquin). Where indicated, proliferation was assessed by pre-labeling cells with CellTrace Violet (Thermo Fisher Scientific) prior to stimulation following manufacturer’s recommendations.

For pharmacological inhibition experiments, inhibitors were added either at the time of stimulation or on day five or six after stimulation for 24 h or 6 h, as specified in the corresponding figure legends. mTORC1 signaling was inhibited using rapamycin (1 µM; Enzo LifeSciences). PI3K/AKT signaling pathway was inhibited using idelalisib (1 µM; Bio-Connect), MK2206 (1 µM; Bio-Connect), or afuresertib (1 µM; Bio-Connect). For lysosomal inhibition, bafilomycin A1 (20 nM; Bio-Connect) was added for 6 h either concurrently with stimulation or after 6 days of stimulation, after which cells were harvested for downstream analyses. For glutamine deprivation experiments, cells were cultured in RPMI 1640 medium without L-glutamine (Gibco) and supplemented as described above.

#### Flow cytometry antibody staining

Cells were stained with eBioscience™ Fixable Viability Dye eFluor™ 780 (Thermo Fisher Scientific) together with fluorochrome-conjugated antibodies against surface proteins surface to enable exclusion of dead cells and identification of immune cell subsets, respectively. All reagents used are listed in the Key Resources Table. Surface staining was performed for 20 min at 4°C in FACS buffer consisting of phosphate-buffered saline (PBS) supplemented with 0.5% fetal bovine serum (Takara) and 0.02% sodium azide.

For intracellular protein staining, surface-stained cells were fixed and permeabilized using the BD Cytofix/Cytoperm™ Fixation/Permeabilization Kit (BD Biosciences) for 20 min at 4°C. Cells were subsequently washed with 1× BD Perm/Wash™ buffer and stained for intracellular targets for 30 min in 1× BD Perm/Wash™ buffer. All antibodies used for intracellular staining are listed in the Key Resources Table. Phosphorylated ribosomal protein S6 was detected using anti-phospho-S6 ribosomal protein (Ser240/244) antibody (Cell Signaling Technology) followed by staining with a goat-anti-rabbit secondary antibody (Southern Biotech). For intracellular cytokine analysis, cells were incubated with Brefeldin A and GolgiStop™ (BD Biosciences) for 4 h at 37°C and 5% CO₂ prior to harvest following the indicated stimulation period. For assessment of degranulation, anti-CD107a antibody was added at the start of this 4 h incubation period. Cells were then harvested and stained for surface markers followed by intracellular cytokine staining using the fixation/permeabilization protocol described above.

#### Quantification of GLUT1 and ASCT2 surface expression

Cell surface expression of the nutrient transporters GLUT1 (glucose transporter 1; *SLC2A1*) and ASCT2 (glutamine transporter; *SLC1A5*) was assessed using ligand-based detection reagents recognizing the extracellular domains of these transporters (Metafora Biosystems). For GLUT1 detection, cells were incubated for 20 min at 37°C with a recombinant viral receptor-binding domain (RBD) probe specific for human GLUT1 fused to GFP (GLUT1.RBD). For ASCT2 detection, cells were incubated for 20 min at 37°C with an RBD probe specific for human ASCT2 fused to a recombinant mouse Fc of IgG (ASCT2.RBD), followed by washing and incubation with a fluorochrome-conjugated anti-mouse secondary antibody (Invitrogen) for 20 min at 4°C. Afterwards cells were washed and subjected to conventional surface staining with viability dye eFluor™ 780 and antibodies as described above.

#### Mitochondrial mass quantification

Mitochondrial mass was assessed by flow cytometry using MitoTracker™ Green FM (Thermo Fisher Scientific). Cells were incubated with 5 nM MitoTracker™ Green FM in 1× Hank’s Balanced Salt Solution (HBSS) containing calcium and magnesium (Gibco) for 15 min at 37°C in a humidified incubator with 5% CO₂. Following staining, cells were washed with cold HBSS and subsequently stained with viability dye eFluor™ 780 and antibodies as described above.

#### Flow cytometry analysis

Flow cytometry data were acquired using BD FACSCanto™ II, BD FACSymphony™ A1, or BD LSRFortessa™ cytometers (BD Biosciences), operated with BD FACSDiva™ software. Instrument choice depended on panel configuration and fluorochrome requirements. Compensation, gating, and downstream data analysis were performed using FlowJo software v10.10.1 (BD Biosciences).

#### Measurement of protein synthesis and cellular energetics through noncanonical amino acid tagging (CENCAT)

PBMCs were stimulated for the indicated time. Cellular energetics through noncanonical amino acid tagging (CENCAT) was performed as previously described by Vrieling et al.,^55^. Specifically, cells were pre-incubated with metabolic inhibitors for 15 min at 37°C in a humidified incubator with 5% CO₂. Treatments included vehicle control (DMSO), 2-deoxy-D-glucose (2-DG; 0.1 M; Merck), oligomycin (1 µM; Sigma-Aldrich), or the combination of 2-DG and oligomycin. Subsequently, the alkyne-bearing noncanonical amino acid β-ethynylserine^56^ (βES; 500 µM), provided as βES-HCl by the laboratory of Dr. Kimberley Bonger, was added for 30 min at 37°C in the continued presence of the respective inhibitors to label nascent protein synthesis. Following incubation with βES, cells were washed with PBS and stained with viability dye eFluor™ 780 together with antibodies against surface proteins for 20 min at 4°C. Cells were then fixed and permeabilized using the BD Cytofix/Cytoperm™ Fixation/Permeabilization Kit for 20 min at 4°C. Incorporated βES was fluorescently labeled by copper-catalyzed azide–alkyne cycloaddition (CuAAC; click chemistry). Click reactions were performed in click reaction mix containing: 1× BD Perm/Wash™ buffer with CuSO₄ (0.5 mM), sodium ascorbate (10 mM), THPTA (2 mM; Merck), aminoguanidine (10 mM; Cayman Chemical), and AFDye™ 405 Azide Plus (0.5 µM; Vector Laboratories). Cells were incubated in click reaction mixture for 30 min at room temperature protected from light, washed with 1× BD Perm/Wash™ buffer, and resuspended in FACS buffer for analysis by Flow Cytometry.

Metabolic dependencies and energetic capacities were calculated from changes in mean fluorescence intensity (MFI) of βES-labeled nascent protein synthesis in response to metabolic inhibition. MFI of the control treatment were used as a measure of protein synthesis.

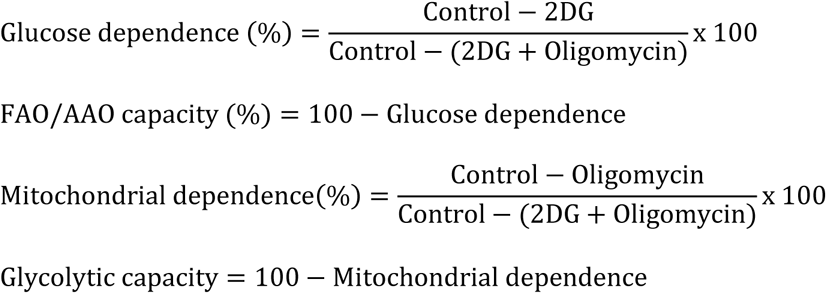

#### Extracellular flux analysis

Extracellular flux analysis was performed as described in detail by Goedhart and Simon-Molas^57^. PBMCs from HD and CLL patients were cultured with or without stimulation for two days. After culture, T cells were isolated from PBMC by initial depletion of CD19+ B cells using anti-CD19 beads for magnetic-activated cell sorting (MACS; Miltenyi Biotec), followed by negative selection of T cells using the EasySep™ Human T Cell Enrichment Kit (STEMCELL Technologies) according to manufacturer’s instructions. Enriched fractions were analyzed by FACS to ensure sufficient T cell purity. Cells were counted manually in a Neubauer chamber using trypan blue staining (ref commercial brand). 50 × 10^3 isolated T cells were seeded in XF HS PDL Miniplates (Agilent Technologies) in RPMI-1640 Medium (Merck), supplemented with 25 mM glucose, 2 mM L-glutamine, and 1 mM sodium pyruvate (pH 7.4, filter sterilized). Plates were centrifuged at 1700 rpm for 5 minutes with brake 0 to immobilize T cells at the bottom of the wells, the correct formation of the cell monolayer was verified by microscope. Plates were incubated in a non-CO₂ incubator at 37°C for 45 min prior to measurement. XF cartridges were hydrated overnight with XF calibrant (Agilent Technologies) following manufacturer’s recommendations. Oxygen consumption rate (OCR) and extracellular acidification rate (ECAR) were measured using an Agilent Seahorse XF HS Mini Extracellular Flux Analyzer with the T cell metabolic profiling kit (Agilent Technologies). The assay consisted of sequential injections of oligomycin (1.5 µM), BAM15 (2.5 µM), and rotenone/antimycin A (0.5 µM each). Basal OCR was calculated from the third measurement prior to oligomycin injection minus non-mitochondrial respiration, defined as the OCR remaining after rotenone and antimycin A treatment. Maximal OCR was calculated as the highest OCR measured following BAM15 injection minus non-mitochondrial respiration. SRC was calculated as maximal OCR minus basal OCR. Basal ECAR was calculated from the third measurements prior to oligomycin injection.

To assess mitochondrial function independently of plasma membrane transport processes and cytoplasmic metabolite handling, extracellular flux analysis was performed on permeabilized cells as described by Goedhart and Simon-Molas 2025. T cells were stimulated and isolated from PBMCs as described above. 50 × 10^3 cells per well were plated in XF HS PDL Miniplates (Agilent Technologies) in 1× mitochondrial assay solution (MAS) consisting of distilled water supplemented with 200 mM mannitol, 70 mM sucrose, 10 mM KH₂PO₄, 5 mM MgCl₂, 1 mM EGTA, and 2 mM HEPES (pH 7.4, filter sterilized). Plates were centrifuged as described above. Assay medium was supplemented with ADP (4 mM), bovine serum albumin (0.2%), pyruvate (5 mM), malate (5 mM), glutamate (5 mM), palmitoyl-carnitine (40 µM), and succinate (5 mM). Selective permeabilization of the plasma membrane was achieved using recombinant perfringolysin O (4 nM; XF Plasma PMP, Agilent Technologies), which allows substrate access to mitochondria while preserving mitochondrial integrity. XF cartridges were hydrated overnight with XF calibrant following manufacturer’s recommendations. The assay was initiated immediately without incubation in a non-CO₂ incubator. OCR and ECAR were measured using the Seahorse XF HS Mini Analyzer (Agilent Technologies). Mitochondrial function was assessed using a mitochondrial stress test protocol with sequential injections of oligomycin (1.5 µM), FCCP (2.5 µM), and rotenone/antimycin A (0.5 µM each). Instrument settings were adjusted to minimize assay duration (mix: 0.5 min; wait: 0 min; measure: 2 min) and reduce cell detachment. In this permeabilized system, ECAR does not reflect glycolytic activity but was used as a quality control parameter to monitor pH stability (target pH 7.4 ± 0.1). Basal phosphorylating respiration was defined as ADP-stimulated oxygen consumption coupled to ATP synthesis via Complex V. All extracellular flux analysis data were analyzed using the Seahorse Analytics online platform.

#### Confocal imaging of mitochondrial morphology

PBMCs from HD and CLL patients were cultured with or without stimulation as described above. To enrich for T cells in CLL PBMC cultures, samples were depleted of CD19+ B cells using MACS. For visualization of mitochondrial morphology, cells were stained with 12.5 nM MitoTracker™ Orange CMTMRos (Thermo Fisher Scientific) in 1× HBSS containing calcium and magnesium (Gibco) for 15 min at 37°C and 5% CO₂. Cells were subsequently stained with fluorochrome-conjugated antibodies against CD4 and CD8 (both Alexa Fluor™ 594-conjugated) diluted in FACS buffer for 40 min at room temperature. Following staining, cells were seeded onto poly-D-lysine (Sigma-Aldrich) coated coverslips in 24-well plates and allowed to adhere for 30 min. Non-adherent cells were gently removed by aspiration. Cells were then fixed with 4% paraformaldehyde (Merck) for 10 min at room temperature, washed, and counterstained with DAPI (Sigma-Aldrich) for 10 min. Coverslips were mounted onto glass slides using Fluoroshield mounting medium (Sigma-Aldrich), allowed to dry for 15 min in the dark at room temperature, and stored at 4°C overnight. Imaging was performed within 24 h after sample mounting. Confocal images were acquired using a Leica TCS SP8 X DLS confocal microscope (Leica Microsystems) controlled by LAS X equipped with a 63× oil-immersion objective. Fluorophores were excited and detected using the following settings: DAPI (360 nm excitation, 460 nm emission), Alexa Fluor 594 (594 nm excitation, 620 nm emission), and MitoTracker™ Orange (550 nm excitation, 576 nm emission). All acquisition parameters were kept constant across experimental conditions to ensure comparability.

Mitochondrial morphology and network parameters were quantified using Fiji^58^ with the Mitochondria Analyzer macro, as previously described by Chaudhry et al.,^59^. T cells were identified and selected in the images based on CD4 and CD8 staining. The mitochondrial channel (MitoTracker) was isolated for analysis. Mitochondrial segmentation was performed using the 2D Threshold Optimize function of the Mitochondria Analyzer macro. Threshold parameters were optimized using a representative subset of images and subsequently applied uniformly across all images within the same experiment. Binarized images were visually inspected to ensure accurate segmentation of mitochondrial structures. Quantitative analysis of mitochondrial morphology and network features (including fragmentation and connectivity metrics) was performed using the 2D analysis module of Mitochondria Analyzer. All analyses were conducted using identical settings across experimental groups. Elongation and complexity were calculated as:

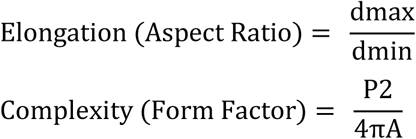

#### Transmission electron microscopy (TEM)

PBMCs from HD and CLL patients were culture with or without stimulation as described above. To enrich for T cells in the CLL PBMC culture, samples were depleted of CD19+ cells using MACS cell separation (Miltenyi Biotec). Cells were the pelleted and immediately fixed in 1% glutaraldehyde, 4% paraformaldehyde (Electron Microscopy Sciences). For embedding samples were post-fixed with 1% osmium tetroxide (OsO4, Electron microscopy sciences) for one hour. Subsequently, samples were dehydrated in an alcohol series from 70% until 100% and last steps in 1,2-propylene oxide (Sigma-Aldrich), then 1:1 ratio 1,2-propylene oxide:epon (LX-112 resin Ladd research) and finally embedded in full epon. Ultrathin (60 nm) epon sections were cut using a diamond (Diatome) knife on a Leica Ultracut UC7 microtome, and collected on 100 mesh cupper grid (Aurion), and counterstained with uranyl acetate and lead citrate (Electron Microscopy Sciences).

Samples were examined using a Talos L120C transmission electron microscope (Thermo Fisher Scientific, Eindhoven, The Netherlands), and images were acquired with a BM-CETA camera using Velox software (Thermo Fisher Scientific, Eindhoven, The Netherlands) or a Jeol JEM-1400 Flash transmission electron microscope equipped with a Xarosa camera and Radius software (EMSIS).

#### Metabolomics

PBMCs from HD and CLL patients were stimulated for two days as described above. After culture, T cells were isolated from PBMC by initial depletion of CD19+ B cells using MACS, followed by fluorescence-activated cell sorting (FACS) using the BD FACSAria II Cell Sorter based on viability staining and CD4 and CD8 expression. Sorted CD4 and CD8 T cells were pelleted at 4°C, washed twice with ice-cold NaCl 0.9% and stored at -80°C prior to analysis. 2 × 10^6^ cells were pelleted per sample. Metabolomics analysis was performed as previously described by Schomakers et al.,^60^, with minor adjustments. The following amounts of internal standard dissolved in water were added to washed cell pellets in a 2 mL tube: adenosine-15N5-monophosphate (5 nmol), adenosine-15N5-triphosphate (5 nmol), D4-alanine (0.5 nmol), D7-arginine (0.5 nmol), D3-aspartic acid (0.5 nmol), D3-carnitine (0.5 nmol), D4-citric acid (0.5 nmol), 13C1-citrulline (0.5 nmol), 13C6-fructose-1,6-diphosphate (1 nmol), guanosine-15N5-monophosphate (5 nmol), guanosine-15N5-triphosphate (5 nmol), 13C6-glucose (10 nmol), 13C6-glucose-6-phosphate (1 nmol), D3-glutamic acid (0.5 nmol), D5-glutamine (0.5 nmol), D5-glutathione (1 nmol), 13C6-isoleucine (0.5 nmol), D3-lactic acid (1 nmol), D3-leucine (0.5 nmol), D4-lysine (0.5 nmol), D3-methionine (0.5 nmol), D6-ornithine (0.5 nmol), D5-phenylalanine (0.5 nmol), D7-proline (0.5 nmol), 13C3-pyruvate (0.5 nmol), D3-serine (0.5 nmol), D6-succinic acid (0.5 nmol), D5-tryptophan (0.5 nmol), D4-tyrosine (0.5 nmol), D8-valine (0.5 nmol). Subsequently, solvents were added to achieve a total volume of 500 µL water, 500 µL methanol and 1 mL chloroform. After thorough mixing, samples were centrifuged for 10 min at 20,000 g. The top layer, containing the polar phase, was transferred to a new 1.5 mL tube and dried using a miVac vacuum concentrator at 60°C. Dried samples were reconstituted in 100 µL 6:4 (v/v) methanol:water. Metabolites were analyzed using a Waters Acquity ultra-high performance liquid chromatography system coupled to a Bruker Impact II™ Ultra-High Resolution Qq-Time-Of-Flight mass spectrometer. Samples were kept at 12°C during analysis and 5 µL of each sample was injected. Chromatographic separation was achieved using a Merck Millipore SeQuant ZIC-cHILIC column (PEEK 100 x 2.1 mm, 3 µm particle size). Column temperature was held at 30°C. Mobile phase consisted of (A) 1:9 (v/v) acetonitrile:water and (B) 9:1 (v/v) acetonitrile:water, both containing 5 mmol/L ammonium acetate. Using a flow rate of 0.25 mL/min, the LC gradient consisted of: 100% B for 0-2 min, reach 54% B at 13.5 min, reach 0% B at 13.51 min, 0% B for 13.51-19 min, reach 100% B at 19.01 min, 100% B for 19.01-19.5 min. Equilibrate column by increasing flow rate to 0.4 mL/min at 100% B for 19.5-21 min. MS data were acquired using negative and positive ionization in full scan mode over the range of m/z 50-1200. Data were analyzed using Bruker TASQ software version 2.1.22.3. The reported metabolite intensities were normalized to internal standards with comparable retention times and response in the MS and to total metabolite pool within each sample. Metabolite identification was based on a combination of accurate mass, (relative) retention times and fragmentation spectra, compared to the analysis of a library of standards. Principal component analysis was performed using the Statistical Analysis function of MetaboAnalyst v5.0^61^ with auto-scaling of data. For heatmap representation, metabolite abundances were standardized by Z-score scaling.

#### 13C glucose and 13C glutamine tracing

PBMCs from HD and CLL patients were stimulated for two days as described above. After culture, T cells were isolated from PBMC by initial depletion of CD19+ B cells using MACS, followed by FACS using the BD FACSAria II Cell Sorter based on viability staining and CD4 and CD8 expression. A minimum of 0,5 × 10^6^ cells were pelleted per sample. Sorted CD4 and CD8 T cells were washed with DMEM without glucose, glutamine and phenol red (Thermo Fisher Scientific) (hereafter, blank medium) and cultured together at 1 × 10^6 cells/mL in blank DMEM supplemented with the corresponding nutrients for isotope labelling. For glucose isotope labelling, 11mM D-Glucose-^13^C_6_ (Sigma-Aldrich) and 2 mM unlabeled glutamine were added. For glutamine isotope labelling, 2mM Glutamine-^13^C_5_ (Cambridge Isotope Laboratories) and 11mM unlabeled glucose were added. T cells were incubated in these media at regular culture conditions for 4 h and subsequently pelleted at 4°C, washed twice with ice-cold NaCl 0.9% and stored at -80°C prior to analysis.

Samples were analyzed using a ZIC-cHILIC-based semi-targeted LC-MS platform following a liquid–liquid extraction, as described above for metabolomics analysis. In this case, Adenosine-^15^N_5_-monophosphate (5 nmol) was used as internal standard. Data were analyzed using Bruker TASQ software version 2.1.22.3. The relative isotope contribution was calculated using IsoCorrectoR Release 3.13^62^. Mean enrichment was calculated by adding the relative contribution of all isotopologues except for M+ 0 and represents the ^13^C-labeled fraction of a given metabolite relative to the total amount of that metabolite within a sample.

#### Intracellular glutamine and malate measurements

PBMCs from HD and CLL patients were cultured with or without stimulation as described above for two days. After culture, T cells were isolated from PBMC by initial depletion of CD19+ B cells using MACS, followed by negative selection of T cells using the EasySep™ Human T Cell Enrichment Kit according the manufacturer’s instructions (malate assessment) or by FACS using the BD FACSAria II Cell Sorter (glutamine assessment). Following isolation, T cells were counted, pelleted, and resuspended in 30 μL PBS per 4 × 10⁴ cells. Samples were lysed by adding 15 μL of Inactivation Solution I per 30 μL initial volume and incubating for 5 min with agitation, followed by 15 μL of Neutralization Buffer per 30 μL initial volume with 1 min of agitation. Final cellular density in lysates was 666 cells/µl. Lysates were stored at −80°C prior to analysis.

Intracellular glutamine levels were measured using the Glutamine/Glutamate-Glo™ Assay (Promega) according to the manufacturer’s instructions. The assay quantifies glutamine indirectly through its enzymatic conversion into glutamate and the subsequent conversion of glutamate into α-ketoglutarate with production of NADH. This reaction is coupled to a NADPH-dependent reductase reaction of a pro-luciferin substrate intro luciferin, which emits light. For measurement, two 25 μL aliquots per sample were transferred into a white 96-well luminometer plate. To enable glutamine conversion, 25 μL glutaminase in glutaminase buffer was added to one set of wells, while 25 μL glutaminase buffer alone was added to paired control wells. Plates were shaken for 30–60 s and incubated for 40 min at room temperature. Subsequently, 50 μL glutamate detection reagent containing glutamate dehydrogenase and reductase was added to all wells, followed by shaking for 30–60 s and incubation for 60 min at room temperature prior to luminescence measurement. Malate levels were measured using the Malate-Glo™ Assay (Promega) according to the manufacturer’s instructions. This assay is based on the coupled reaction of malate dehydrogenase, which produces NADH, with a reductase that converts pro-luciferin into luciferin. 50 μL of each sample were transferred into a white 96-well luminometer plate, followed by addition of 50 μL malate detection reagent containing the enzymes. Plates were shaken for 30–60 s and incubated for 60 min at room temperature prior to luminescence measurement. For both assays, luminescence was measured using a BioTek luminometer

#### RNAseq

PBMCs from HD and CLL patients were stimulated for two days as described above. After culture, T cells were isolated from PBMC by initial depletion of CD19+ B cells using MACS (Miltenyi Biotec), followed by FACS using the BD FACSAria II Cell Sorter based on viability staining and CD4 and CD8 expression. A minimum of 1 × 10^6^ cells were pelleted per sample, lysed and total RNA was isolated using the RNeasy mini kit (74106, Qiagen) according to the manufacturer’s protocol. RNA quality was assessed using a Fragment Analyzer and libraries were generated using the NEBNext Ultra II Directional RNA library prep kit for Illumina. Samples were barcoded and sequenced using a NovaSeq6000 (PE 150 bp). Quality of the sequencing data was assessed using FastQC. Raw FastQ files were aligned to the GRCh38 human genome using STAR v2.7.9a and the transcriptome.out.bam files were used as input for RSEM’s calculate expression (v1.3.3) to generate expected counts^63^. Normalized counts were generated with DEseq2 and plotted with pheatmap function, significant differential expression was determined at FDR < 0.05 ^64^. The distribution of differentially expressed genes between HD and CLL T cells was assessed within the Hallmark gene sets MTORC1 SIGNALING, MYC TARGETS V1, and MYC TARGETS V2^65^. Raw data has been deposited in the European Genome-phenome Archive (EGA; EGAD50000001368).

#### Statistical analysis

Data were analyzed using GraphPad Prism software v10.1.2. Data are presented as mean ± standard error of the mean and were analyzed, as indicated in the figure legends with two-tailed Student’s t test (paired according to experimental setting), simple linear regression, ordinary two-way or one-way analysis of variance (ANOVA) followed by Tukey multiple-comparisons test or Šídák multiple comparisons tests with matched data. P value < 0.05 was considered statistically significant.

