## Supplementary Tables for "Glutamine-driven mTORC1 activity enforces glycolytic bias, hyperactivation and aberrant proliferation in T cells in chronic B-cell leukemia"

**Table S1**

| Patient ID | Sex | Age | IgVH mutation status | Rai stage | Leucocytes (WBC) 10e9/L | % CD5+CD19+ (CLL cells) on total lympho | % CD3+ on total lympho | Treated at sample date | Treatment |
|---|---|---|---|---|---|---|---|---|---|
| CLL01 | Female | 81 | N/A | 0 | 20.9 | 87 | 7.2 | No | |
| CLL02 | Male | 59 | Mutated | III | 58.7 | 93 | 4.4 | No | |
| CLL03 | Male | 75 | Unmutated | I | 127 | 93 | 12.3 | No | |
| CLL04 | Female | 69 | N/A | 0 | 49 | 90.35 | 4.05 | No | |
| CLL05 | Female | 62 | Mutated | 0 | 173 | 99.9 | 7.4 | Yes | Chlorambucil (Leukeran) |
| CLL06 | Female | 55 | N/A | IV | 16.3 | 94.8 | 23.1 | No | CHOP (cyclophosphamide, doxorubicin, vincristine, prednisolone) |
| CLL07 | Male | 70 | Mutated | 0 | 67.92 | 94.14 | 4.19 | No | |
| CLL08 | Female | 68 | Mutated | 0 | 91.81 | 91.67 | 7.34 | No | |
| CLL09 | Female | 84 | Mutated | 0 | 63.77 | 92.65 | 5.92 | No | |
| CLL10 | Female | 81 | Unmutated | 0 | 25.87 | 85.71 | 10.62 | No | |
| CLL11 | Male | 64 | Mutated | 0 | 13.93 | 74.89 | 20.94 | No | |
| CLL12 | Female | 69 | Mutated | 0 | 24.9 | 89.93 | 7.54 | No | |
| CLL13 | Female | 76 | Mutated | I | 46.16 | 92.62 | 6.14 | No | |
| CLL14 | Female | 79 | Mutated | 0 | 47.11 | 70.63 | 11.71 | No | |
| CLL15 | Male | 85 | Mutated | 0 | 50.41 | 88.89 | 9.44 | No | |
| CLL16 | Female | 54 | Mutated | I | 76.84 | 94.52 | 4.12 | No | |
| CLL17 | Female | 82 | Mutated | II | 75.7 | 95.4 | 3.4 | Yes | XELODA (capecitabine) |
| CLL18 | Male | 71 | Mutated | 0 | 44.96 | 91.16 | 7.23 | No | |
| CLL19 | Male | 78 | Mutated | 0 | 85.32 | 92.84 | 5.13 | No | |
| CLL20 | Male | 70 | Mutated | 0 | 125.34 | 96.05 | 3.57 | No | |
| CLL21 | Female | 91 | Unmutated | II | 56.19 | 91.48 | 6.57 | No | |
| CLL22 | Male | 71 | Mutated | I | 145.56 | 92.34 | 6.07 | No | |
| CLL23 | Female | 63 | Mutated | 0 | 170.06 | 96.07 | 3.35 | No | |
| CLL24 | Male | 63 | N/A | N/A | 66.15 | 86.78 | 11.55 | No | |
| CLL25 | Male | 67 | Mutated | 0 | 161.68 | 93.19 | 2.5 | No | |
| CLL26 | Female | 74 | Mutated | 0 | 21.04 | 91.25 | 7.07 | No | |
| CLL27 | Male | 70 | Mutated | 0 | 43.26 | 87.27 | 11.08 | No | |
| CLL28 | Female | 73 | Mutated | I | 36.68 | 90.43 | 8.05 | No | |
| CLL29 | Male | 73 | Mutated | 0 | 52.78 | 92.33 | 5.52 | No | |
| CLL30 | Male | 57 | Mutated | II | 97.79 | 97.79 | 1.8 | No | |
| CLL31 | Male | 66 | Mutated | I | 73.41 | 91.98 | 5.94 | No | |
| CLL32 | Female | 63 | Mutated | 0 | 80 | 94 | 4.2 | Yes | FCR (fludarabine + cyclophosphamide + rituximab) |
| CLL33 | Male | 78 | Mutated | 0 | 51.18 | 90.94 | 7 | No | |
| CLL34 | Female | 81 | Mutated | 0 | 41.91 | 85.75 | 10.69 | No | |
| CLL35 | Male | 60 | Mutated | II | 17.81 | 85.19 | 11.25 | No | |
| CLL36 | Female | 75 | Mutated | III | 292.68 | 98.48 | 1.31 | Yes | Corticosteroids |
| CLL37 | Female | 66 | Mutated | 0 | 13.75 | 79.44 | 17.27 | No | |
| CLL38 | Male | 72 | Mutated | I | 54.91 | 95.74 | 3.35 | No | |
| CLL39 | Female | 60 | N/A | I | 40.24 | 87.74 | 10.59 | No | |
| CLL40 | Male | 61 | Mutated | III | 200.68 | 94.82 | 4.43 | No | |
| CLL41 | Male | 51 | Mutated | I | 167.31 | 96.05 | 3.06 | No | |
| CLL42 | Male | 66 | Unmutated | I | 58.33 | 92.13 | 5.22 | No | |
| CLL43 | Male | 73 | N/A | 0 | 43.27 | 87.47 | 10.97 | No | |
| CLL44 | Female | 71 | N/A | 0 | 210.69 | 92.28 | 6.03 | No | |
| CLL45 | Male | 74 | N/A | II | 92.47 | 94.03 | 5.82 | No | |
| CLL46 | Female | 83 | Mutated | I | 38.2 | 86.86 | 10.74 | No | |
| CLL47 | Male | 85 | Mutated | II | 125.38 | 98.46 | 0.96 | No | |
| CLL48 | Male | 72 | N/A | I | 42.52 | 93.08 | 5.45 | No | |
| CLL49 | Male | 78 | N/A | 0 | 112 | 92.8 | 1.9 | No | |
| CLL50 | Female | 78 | Unmutated | N/A | 73.17 | 91.32 | 7.23 | No | |
| CLL51 | Female | 78 | Mutated | I | 91.17 | 93.59 | 5.29 | No | |
| CLL52 | Male | 69 | N/A | 0 | 61 | 88.9 | 6 | No | |
| CLL53 | Female | 63 | N/A | N/A | 77.9 | 91.83 | 6.78 | No | |

**Table S2**

| Patient ID | Sex | Age | Mutation status |
|---|---|---|---|
| CLL01 | Male | 68 | Mutated |
| CLL02 | Male | 63 | Mutated |
| CLL03 | Male | 73 | Unmutated |
| CLL04 | Female | 66 | Unmutated |
| CLL05 | Male | 60 | Unmutated |
| CLL06 | Male | 63 | Mutated |
| CLL07 | Male | 46 | Unmutated |
| CLL08 | Male | 64 | Mutated |
| CLL09 | Male | 55 | Mutated |
| CLL10 | Male | 71 | Mutated |
| CLL11 | Male | 54 | Mutated |
| CLL12 | Male | 50 | Unmutated |
| CLL13 | Male | 50 | Unmutated |
| CLL14 | Male | 69 | Unmutated |
| CLL15 | Female | 66 | Mutated |
| CLL16 | Male | 57 | Unmutated |
| CLL17 | Female | 65 | Mutated |
| CLL18 | Male | 63 | Mutated |
| CLL19 | Female | 69 | Mutated |
| CLL20 | Male | 58 | Mutated |
| CLL21 | Female | 58 | Unmutated |
| CLL22 | Male | 59 | N/A |
| CLL23 | Female | 73 | Unmutated |
| CLL24 | Male | 73 | Unmutated |
| CLL25 | Male | 52 | Mutated |
| CLL26 | Female | 52 | Unmutated |
| CLL27 | Male | 67 | Unmutated |
| CLL28 | Male | 47 | Mutated |
| CLL29 | Male | 72 | Unmutated |
| CLL30 | Male | 69 | Unmutated |
| CLL31 | Male | 83 | Mutated |
| CLL32 | Male | 68 | Mutated |
| CLL33 | Female | 55 | Unmutated |
| CLL34 | Male | 46 | Unmutated |
| CLL35 | Female | 63 | Unmutated |
| CLL36 | Male | 65 | Unmutated |
| CLL37 | Male | 61 | Mutated |

**Table S3**

| ID | Sex | Age |
|---|---|---|
| HD01 | Male | 1900 |
| HD02 | Male | 1900 |
| HD03 | Male | 1900 |
| HD04 | Male | 1900 |
| HD05 | Female | 1900 |
| HD06 | Male | 1900 |
| HD07 | Female | 1900 |
| HD08 | Male | 1900 |
| HD09 | Male | 1900 |
| HD10 | Female | 1900 |
| HD11 | Female | 1900 |
| HD12 | Male | 1900 |
| HD13 | Male | 1900 |
| HD14 | Male | 1900 |
| HD15 | Male | 1900 |
| HD16 | Female | 1900 |
| HD17 | Male | 1900 |
| HD18 | Female | 1900 |
| HD19 | Female | 1900 |
| HD20 | Female | 1900 |
| HD21 | Female | 1900 |
| HD22 | Female | 1900 |
| HD23 | Female | 1900 |
| HD24 | Male | 1900 |
| HD25 | Female | 1900 |
| HD26 | Male | 2024 |
| HD27 | Female | 2024 |
| HD28 | Female | 2024 |
| HD29 | Male | 2024 |
| HD30 | Female | 2024 |
| HD31 | Male | 2025 |
| HD32 | Male | 2025 |
| HD33 | Female | 2025 |
| HD34 | Female | 2025 |
| HD35 | Female | 2025 |
| HD36 | Female | 2025 |
| HD37 | Male | 2025 |
| HD38 | Female | 2025 |
| HD39 | Female | 2025 |
| HD40 | Female | 2025 |
| HD41 | Female | 2025 |
| HD42 | Male | 2025 |
