## Supplementary material for "Glutamine-driven mTORC1 activity enforces glycolytic bias, hyperactivation and aberrant proliferation in T cells in chronic B-cell leukemia": Key resources table

| Reagent or Resource | Source | Identifier |
|---|---|---|
| **Antibodies** | | |
| anti-human CD27 (clone L128), BUV395 | BD Biosciences | 563816 |
| anti-human CD25 (clone M-A251), PE | BD Biosciences | 555432 |
| anti-human CD25 (clone M-A251), BV786 | BD Biosciences | 563701 |
| anti-human CD69 (clone L78), PE | BD Biosciences | 341652 |
| anti-human CD71 (clone OKT-09), FITC | eBioscience | 11-0719-42 |
| anti-human CD4 (clone RPA-T4), FITC | BD Biosciences | 555346 |
| anti-human CD4 (clone RPA-T4), BV605 | BD Biosciences | 562658 |
| anti-human CD4 (clone RPA-T4), BV650 | BioLegend | 300536 |
| anti-human CD45RA (clone HI100), BV605 | BD Biosciences | 562886 |
| anti-human CD8 (clone RPA-T8), PE-Cy7 | eBioscience | 25-0088-42 |
| anti-human CD8 (clone RPA-T8), BV786 | BD Biosciences | 563823 |
| anti-human CD8 (clone RPA-T8), V450 | BD Biosciences | 560347 |
| anti-human Phospho-S6 Ribosomal Protein Ser240/244 (polyclonal) | Cell Signaling Technology | 2215 |
| anti-human CD107a (clone H4A3), PE-Cy7 | BD Biosciences | 561348 |
| anti-human Granzyme B (clone GB11), Alexa Fluor 700 | BD Biosciences | 560213 |
| anti-human IFNγ (clone B27), BV421 | BD Biosciences | 562988 |
| anti-human TNFα (clone MAB11), BV650 | BD Biosciences | 563418 |
| anti-human IL-2 (clone MQ1-17H12), PE/Dazzle 594 | BioLegend | 500343 |
| Phosflow™ Mouse Anti-AKT T308 (clone J1-223.371), PE | BD Biosciences | 558275 |
| Phospho-AKT1 Ser473 (clone SDRNR), APC | eBioscience | 17-9715-42 |
| Goat anti-rabbit IgG (polyclonal), PE | SouthernBiotech | 4010-09S |
| Goat anti-mouse IgG (polyclonal), Alexa Fluor 647 | Invitrogen | A21235 |
| **Biological Samples** | | |
| PBMC from healthy donor | Sanquin Blood Bank, The Netherlands | N/A |
| PBMC from CLL patients | B-cell Malignancies Biobank, Amsterdam UMC, The Netherlands | N/A |
| PBMC from CLL patients | University Hospital of Cologne | N/A |
| **Chemicals, Peptides, and Recombinant Proteins** | | |
| RPMI 1640 + L-glutamine | Thermo Fisher Scientific | 22400089 |
| RPMI 1640 Medium, no glutamine | Thermo Fisher Scientific | 21870076 |
| RPMI-1640 Medium | Merck | R1383 |
| HBSS (1X), calcium, magnesium, phenol red | Thermo Fisher Scientific | 15420614 |
| DMEM, no glucose, no glutamine, no phenol red (blank medium) | Thermo Fisher Scientific | A1443001 |
| eBioscience™ Fixable Viability Dye eFluor™ 780 | Thermo Fisher Scientific | 65-0865-14 |
| CellTrace Violet | Thermo Fisher Scientific | C34557 |
| anti-CD3 clone 1XE, purified | Sanquin | N/A |
| anti-CD28 clone 15E8, purified | Sanquin | N/A |
| Rapamycin | Enzo Life Sciences | BML-A275 |
| Idelalisib | Bio-Connect | S2226 |
| MK2206 | Bio-Connect | HY-10358 |
| Afuresertib | Bio-Connect | HY-15727 |
| Bafilomycin A1 | Bio-Connect | 88899-55-2 |
| GolgiStop™ | BD Biosciences | 554724 |
| Brefeldin A | Sigma-Aldrich | B7651 |
| Cytofix/Cytoperm™ Fixation/Permeabilization Kit | BD Biosciences | 554714 |
| MitoTracker™ Green FM | Thermo Fisher Scientific | M7514 |
| MitoTracker Orange CMTMRos | Thermo Fisher Scientific | M7510 |
| Fetal bovine serum for cell culture (tetracycline-free) | Takara | 631101 |
| Penicillin-Streptomycin Solution | Corning | 30-002-CI |
| β-ethynylserine-HCl (βES) | Bonger Lab, Leiden Institute for Chemistry, Chemical Biology & Immunology[56] | N/A |
| 2-deoxy-D-glucose (2-DG) | Merck | D8375 |
| Oligomycin A | Sigma-Aldrich | O4876 |
| BAM15 | Sigma-Aldrich | SML1760 |
| FCCP | Sigma-Aldrich | C2920 |
| Rotenone | Sigma-Aldrich | R8875 |
| Antimycin A | Sigma-Aldrich | A8674 |
| Aminoguanidine | Cayman Chemical | CAY81530 |
| THPTA | Merck | 762342 |
| AZDye 405 Azide Plus | Vector Laboratories | CCT-1474 |
| Ficoll-Paque™ PREMIUM | VWR | 17-5442-03 |
| ADP | Sigma-Aldrich | A5285 |
| Malate | Sigma-Aldrich | 240176 |
| Pyruvate | Sigma-Aldrich | 107360 |
| Glutamate | Sigma-Aldrich | G1251 |
| Palmitoyl-carnitine | Sigma-Aldrich | 61251 |
| Succinate | Sigma-Aldrich | S3674 |
| Paraformaldehyde | Merck | 158127 |
| Poly-D-lysine | Sigma-Aldrich | P6407 |
| DAPI | Sigma-Aldrich | D9542 |
| Fluoroshield mounting medium | Sigma-Aldrich | F6182 |
| 11 mM D-Glucose-13C6 | Sigma-Aldrich | 389374 |
| 2 mM Glutamine-13C5 | Cambridge Isotope Laboratories | CLM-1822-H |
| 1% glutaraldehyde, 4% paraformaldehyde | Electron Microscopy Sciences | 15949-70 |
| Osmium tetroxide ($OsO_4$) | Electron Microscopy Sciences | 20816-12-0 |
| Lead Citrate | Electron Microscopy Sciences | 22410 |
| **Critical Commercial Assays** | | |
| Anti-GLUT-1 RBD conjugated to GFP | Metafora Biosystems | GLUT1.RBD |
| Anti-ASCT2 RBD conjugated to mouse RFc-IgG | Metafora Biosystems | ASCT2.RBD |
| EasySep™ Human T Cell Enrichment Kit | STEMCELL Technologies | 19051 |
| XFp FluxPak (PDL plates) | Agilent Technologies | 103721-100 |
| Seahorse XF Calibrant Solution | Agilent Technologies | 100840-000 |
| Seahorse XF T Cell Metabolic Profiling Kit | Agilent Technologies | 103771-100 |
| Seahorse XF Plasma Membrane Permeabilizer (PMP) | Agilent Technologies | 102504-100 |
| LD Columns | Miltenyi Biotec | 130-042-901 |
| CD19 MicroBeads, human | Miltenyi Biotec | 130-050-301 |
| BD Cytofix/Cytoperm™ Fixation/Permeabilization Kit | BD Biosciences | 554714 |
| Glutamine/Glutamate-Glo™ Assay | Promega | J8021 |
| Malate-Glo™ Assay | Promega | JE9100 |
| **Software and Algorithms** | | |
| FlowJo v10.10 | Tree Star | https://www.flowjo.com/ |
| BD FACSDiva Software | BD Biosciences | https://www.bdbiosciences.com/ |
| Seahorse Wave Desktop Software | Agilent Technologies | https://www.agilent.com/ |
| GraphPad Prism v10 | GraphPad Software | https://www.graphpad.com/ |
| Fiji | Schindelin et al., 2012 | https://imagej.net/software/fiji/ |
| Mitochondria-Analyzer | Chaudhry et al., 2019 | GitHub repository |
| LAS X Software | Leica Microsystems | Leica software portal |
| MetaboAnalyst v5.0 | Pang et al., 2021 | https://www.metaboanalyst.ca/ |
| IsoCorrectoR Release 3.13 | Heinrich et al., 2018 | R package |
| Velox Software | Thermo Fisher Scientific | Thermo Fisher Scientific |
| MetaboAnalyst v5.0 | Pang et al., 2021 | https://www.metaboanalyst.ca/ |
